# High gene flow in Atlantic vent shrimp species with multiple ridge colonizations by ecotypic lineages with contrasting symbioses

**DOI:** 10.64898/2026.01.09.698622

**Authors:** Elodie Portanier, Pierre Methou, Claire Daguin-Thiébaut, Stéphanie Ruault, Anne-Sophie Le Port, Chong Chen, Marjolaine Matabos, Didier Jollivet, Florence Pradillon

## Abstract

Conservation concerns regarding potential mineral extraction near deep-sea hydrothermal vents have emphasised the need for a better understanding of the contemporary connectivity of various vent-endemic species, accounting for the specifics of their life-history traits and habitat requirements. *Rimicaris* shrimps are a major component of vent communities in the Atlantic Ocean, both along the Mid-Atlantic Ridge (MAR) and at the Mid-Cayman Spreading Center (MCSC). Using ddRAD sequencing, we examined the genetic differentiation of three Atlantic *Rimicaris* species to better understand their past and present migration pathways and the possible demographic history that led to their current distribution and ecology. *Rimicaris exoculata* and *R. chacei* inhabit the MAR and exhibit widespread gene flow along the ridge with no areas of restricted connectivity. *R. hybisae* occurs at the MCSC and exhibits almost no genetic difference with its congeneric from the MAR, *R. chacei*, despite the geographic distance between them and their contrasting ecologies and morphologies. Demographic inferences support a scenario of a very recent (less than 2000 generations) separation between MCSC and MAR shrimp populations, followed by a rapid expansion in both regions, possibly triggered by symbiotic development. Analyses further suggest recent or ongoing secondary contacts involving two-way migration, which is more pronounced from the MCSC towards the MAR. All together these results suggest that *R. chacei* and *R. hybisae* represent distinct forms of the same species, which we hereby synonymise as *R. chacei*, highlighting their morphological plasticity in relation to symbiosis.

## 1. Introduction

Deep-sea hydrothermal vents are highly fragmented and unstable habitats hosting endemic and highly specialised species that function as a metacommunity, and where long-distance dispersal of planktonic larvae can mitigate the risk of local extinction (Mullineaux et al., 2018). At the species level, the manner in which populations are connected depends on specific life-history traits, including reproductive effort, larval development and behaviour in the water column, pelagic larval duration and settlement requirements for a given habitat (Adams et al., 2012). For vents, the availability of suitable habitats depends on geological settings that drive the distribution and dynamics of sites and habitat patches (Tunnicliffe & Fowler, 1996). Hence, in a given region, co-distributed vent species may exhibit different population connectivity networks, resulting from their specific combination of biological traits and habitat requirements. This was observed in dynamic systems of the Pacific Ocean either at the fast-spreading East Pacific Rise (EPR), where vent species exhibit different genetic structures with geographical breaks occurring at various places along the ridge (Hurtado et al., 2004; Plouviez et al., 2009), or across back-arc basins of the western Pacific (Poitrimol et al., 2022).

In the Atlantic Ocean, the slower seafloor spreading rates of the Mid-Atlantic Ridge (MAR) and the Mid Cayman Spreading Center (MCSC) result in relatively long-lived vent sites, and populations are more distantly distributed and less ephemeral than those encountered on fast spreading ridges (Beaulieu et al., 2015; Matabos et al., 2025; Van Audenhaege et al., 2024). In addition, vent habitats exhibit marked spatial heterogeneity with a strong depth gradient, as well as contrasting geochemistry (Diehl & Bach, 2020; Konn et al., 2022). This results in different vent communities along the ridge (Desbruyères et al., 2000; Plouviez et al., 2015). Connectivity networks may therefore be particularly sensitive to local (at the vent field scale) disturbances, habitat choice, and self-recruitment, as the rescue effect of larvae travelling from afar may be more easily disrupted than in systems where populations are closer from each other (Mullineaux et al., 2020). In the northern MAR, all known vent fields are currently included in exploration licences for deep seafloor mineral resources, with the exception of the vent fields south of the Azores (Menez Gwen, Lucky Strike and Rainbow) which are included in a Portuguese Marine Natural Reserve. This makes them potentially vulnerable to future mining activities (Menini & Van Dover, 2019). There is thus an urgent need to better understand large-scale connectivity networks, accounting for species with contrasting life history traits, to be able to inform spatial management strategies.

The MAR hydrothermal vents host three main faunal assemblages. Aggregations of the alvinocaridid shrimp *Rimicaris exoculata* occupy the hottest habitats near fluid orifices (Hernández-Ávila et al., 2022; Methou et al., 2022; Segonzac et al., 1993). Patches of the gastropods *Peltospira smaragdina* and *Divia briandi* together with other vent shrimps are found at intermediate temperatures on chimney walls (Cuvelier et al., 2025; Sarrazin et al., 2022), while extensive beds of the *Bathymodiolus* mussels co-occurring with the limpet *Lepetodrilus atlanticus*, are found in cooler habitats (Desbruyères et al., 2000; Sarrazin et al., 2020). Like in dynamic systems such as the EPR, the dominant MAR species display various connectivity patterns. *Bathymodiolus* mussels as well as the gastropods *L. atlanticus* and *P. smaragdina* all exhibit marked genetic structuration along the MAR, with a common region of reduced exchange between 26°N and 36°N (Breusing et al., 2016, 2017; Faure et al., 2009; O’Mullan et al., 2001; Portanier et al. in press). In contrast, species such as the gastropod *Divia briandi* (Yahagi et al., 2019), shrimp *R. exoculata* (Teixeira et al., 2011, 2012) and their commensal dirivultid copepods *Stygiopontius pectinatus* (Gollner et al., 2016) show no genetic structuring across their known geographical distribution.

Demographic histories of species with putative expansion or contraction also influence the present-day spatial patterns of genetic variation. This can promote geographic isolation and genetic barriers when secondary contacts then occur, but also habitat sharing, which can lead to species competition or maladaptation, allowing long-term evolutionary changes, such as speciation or niche partitioning, to occur. Along the MAR, two mussels species, *Bathymodiolus azoricus* and *B. puteoserpentis* display a parapatric distribution along a depth gradient with the presence of a hybrid zone at Broken Spur (29°N) (Breusing et al., 2017; O’Mullan et al., 2001; van der Heijden et al., 2012). Genetic admixture has also been observed in the same region between genetically differentiated populations of the gastropod *P. smaragdina* (Portanier et al., in press), which also unfrequently shares its habitat with another closely-related but chemosymbiotic species *P. gargantua*. This latter situation can likely be explained by specific environmental filters or niche partitioning in certain locations (Chen et al., 2025; Sarrazin et al., 2022).

Alvinocaridid shrimps inhabit deep-water hydrothermal vents and cold seeps around the world, and are generally recognized as good dispersers with high connectivity (Methou, Ogawa, et al., 2024; Thaler et al., 2014; Yahagi et al., 2015; Y. Zhou et al., 2022), although exceptions of restricted regional-scale structures have been reported in *Rimicaris* species from the Mariana Arc (Methou et al., 2025). In the Atlantic, two species stand out due to their highly modified head morphology at the adult stage, where the enlarged cephalothoracic cavity houses a diverse chemosynthetic microbial community involved in the shrimp nutrition: *R. exoculata* along the MAR and *R. hybisae* at MCSC vents, that both form high density assemblages around vent chimneys (Cambon-Bonavita et al., 2021; Nye et al., 2013; Ponsard et al., 2013; Zbinden & Cambon-Bonavita, 2020). Another species: *R. chacei*, which has a much less developed symbiosis and much smaller cephalothoracic cavity, lives sympatrically with *R. exoculata* but occupies a distinct ecological niche (Methou et al., 2020, 2022). Based on mitochondrial markers,

*R. chacei* appears to be genetically very similar to the MCSC species *R. hybisae* (Methou, Guéganton, et al., 2024; Teixeira et al., 2013; Vereshchaka et al., 2015). However, these two species have been named and treated as separate species due to their distinct morphology and ecology (Nye et al., 2012; Streit et al., 2015). Although *R. chacei* has similar microbial communities to *R. exoculata* and *R. hybisae*, the anatomical changes it acquires during metamorphosis to host its symbionts are less pronounced. This has been attributed to differences in how juveniles acquire their symbionts immediately after settlement (Guéganton et al., 2024; Methou, Guéganton, et al., 2024).

The genetic closeness between *R. chacei* and *R. hybisae* could be indicative of past or present-day connections between the MAR and the MCSC, despite them being almost 4000 km apart. Similarly, the genetic structure of *R. exoculata* using both mitochondrial and microsatellite data indicates that shrimp metapopulations are well connected over distances of thousands of kilometres between vent fields of different depths (Teixeira et al., 2011, 2012). However, the use of a limited number of genetic markers in existing studies may not reveal fine-scale genetic structure of the different *Rimicaris* species in the Atlantic. Consequently, it remains unclear whether *R. chacei* from MAR and *R. hybisae* from MCSC represent two geographic species that underwent recent allopatric speciation in their respective regions or whether they are a single species with two distinct morphological ecotypes that still exchange migrants across the Caribbean-Atlantic region. Although existing genetic evidence argues against the ‘two species’ hypothesis, incomplete lineage sorting and low mutation rates in markers used for phylogenetic approaches, can both result in failure to separate semi-isolated species falling in the grey zone of speciation (Roux et al., 2016).

Here, we propose to use genome-wide scans of genetic differentiation in the three Atlantic *Rimicaris* species to better understand past and contemporary migration pathways, and the possible demographic history that led to their current distribution and ecology. These new datasets will help refine the connectivity networks of *Rimicaris* shrimps and identify areas with weaker connections, if any, thus providing key information for mitigating the impact of potential extractive activities near MAR deep-sea vents. The datasets will also be used to determine whether *R. chacei* and *R. hybisae* belong to a single genetic unit with widespread distribution that exhibits ecological plasticity in response to trophic competition. As such, the study will provide insights into how niche partitioning, symbiont acquisition and migration may interact to inform hypotheses regarding the adaptive mechanisms of Atlantic *Rimicaris* shrimps.

## 2. Materials and Methods

### 2.1. Sample collection and DNA extraction

Shrimp samples were collected from eight hydrothermal vent fields on the northern MAR and the MCSC in the Caribbean Sea (see: Figure 1A and Table 1) during nine scientific cruises (Supplementary Tables S1) for *Rimicaris exoculata* (5 localities), *R. chacei* (4 localities) and *R. hybisae* (the 2 known MCSC localities) using either the HOVs *Nautile* and *Shinkai 6500*, or the ROVs *Victor 6000* and *Isis*. After their recovery on board, specimens were either preserved in 96% ethanol or individually frozen at –80°C. DNA was extracted from two to three pleopods of each individual, using 96-wells extraction columns (Nucleospin 96 Tissue, Macherey–Nagel) with negative extraction controls present throughout. The pleopods were digested for 2 hours at 56 °C with 30 µL of Proteinase K (20 mg/mL) under 350 rpm shaking. Lysates were then incubated 20 minutes at ambient temperature with 20 µL of RNAse A (20 mg/mL) and transferred into Nucleospin purification columns and centrifuged at 5600 rpm following manufacturer’s instructions. DNA was then eluted in 100 μL of BE buffer and stored in 96-wells plates at − 20 °C. The quality of DNA samples was checked using electrophoresis on a 0.8% agarose TBE gel stained with ethidium bromide and the quantity was determined using a Qubit fluorimeter (ThermoFisher Scientific).

**Figure 1:**
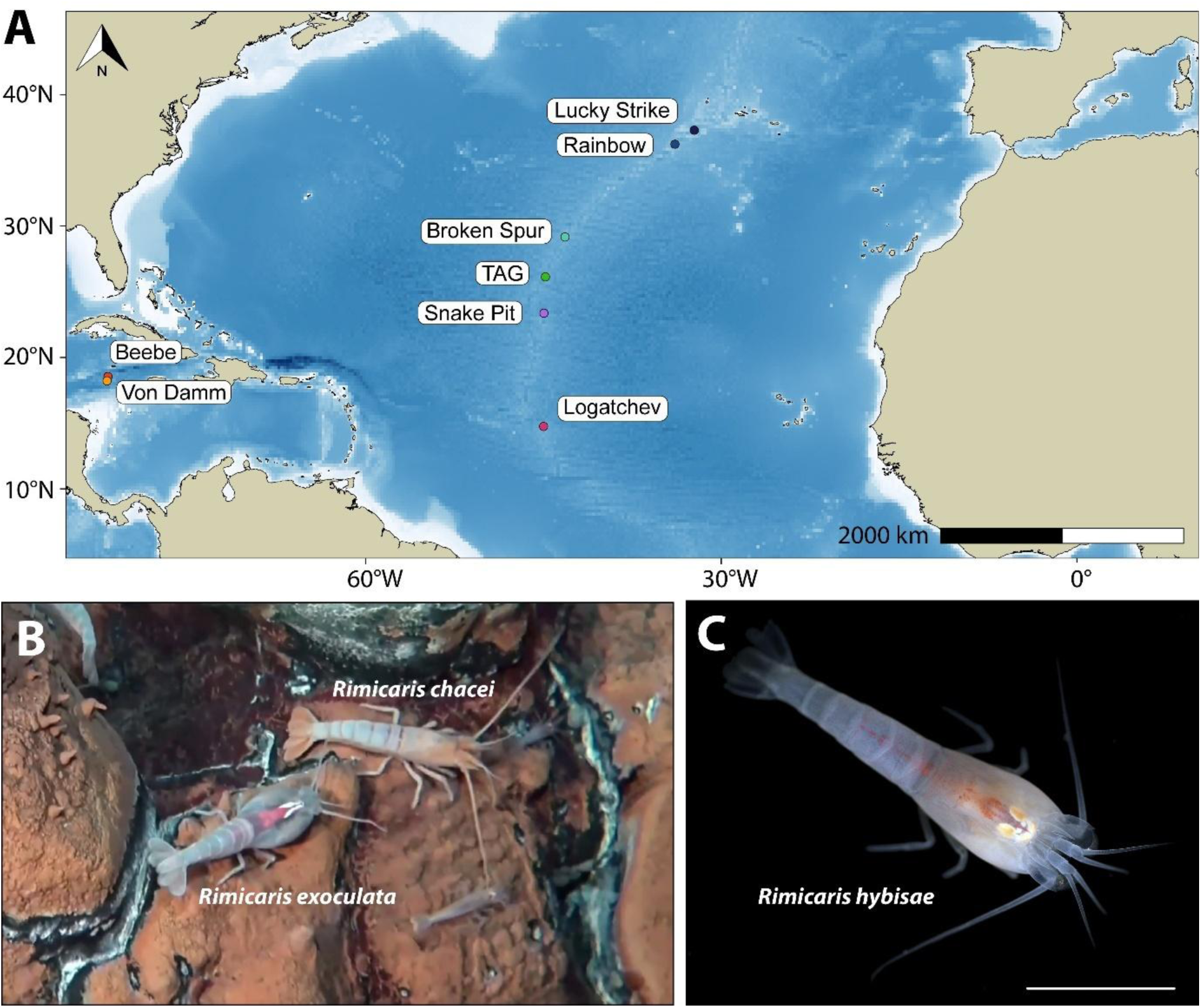
Context of the study. **A.** Map of hydrothermal vent fields sampled in the northern Atlantic and the Caribbean Sea. Same color code for the different vent fields is applied for our genetic analyses **B.** Two individuals of *R. exoculata* (right below) and *R. chacei* (left on top) on a rock close to shimmering fluids at the TAG vent field (Credit: BICOSE 2 expedition; Ifremer). **C.** *R. hybisae* individual from the Von Damm vent field (scale bar = 1mm) (Credit: Dr. Adrian Glover, Natural History Museum UK).

**Table 1:** Numbers of individuals of each species per vent field for which ddRAD data were obtained.

| **Vent field** | ***R. chacei*** | ***R. exoculata*** | ***R. hybisae*** | **Total** |
| --- | --- | --- | --- | --- |
| Lucky Strike | 18 |  |  | **18** |
| Rainbow |  | 24 |  | **24** |
| Broken Spur | 18 | 16 |  | **34** |
| TAG | 67 | 18 |  | **85** |
| Snake Pit | 34 | 20 |  | **54** |
| Logatchev |  | 22 |  | **22** |
| Beebe |  |  | 24 | **24** |
| Von Damm |  |  | 31 | **31** |
| **Total** | **137** | **100** | **55** | **292** |

### 2.2. ddRAD-seq library preparation and sequencing

Individual ddRAD libraries were prepared according to the protocol of (Brelsford et al., 2016), as modified by (Daguin Thiebaut et al., 2021). Approximately 100 ng of DNA was digested for each individual using high-fidelity *SbfI* and *MseI* restriction enzymes (New-England BioLabs). The *SbfI* enzyme was chosen due to its low cutting frequency in genomic DNA, given the very large size of the shrimp’s duplicated genomes (>8 Gb). Ligation with adapters, purification, amplification and pooling were then carried out in our laboratory (see: Daguin-Thiébaut et al. 2021 for details). A 1.5% DF marker K agarose gel cassette was used to select fragments between 300 and 800 pb in a Pippin Prep (SageScience), and the quality of ddRAD libraries was checked on a High Sensitivity DNA Chip using a Bioanalyzer (Agilent^TM^). The pooled libraries were then paired-end sequenced (150 bp) in two steps. First, a subset of 6 individual libraries were pooled and sequenced on a NovaSeq 6000 Illumina sequencer to check the library quality (n = 6 individuals, Genoscope, Centre National de Séquençage, Evry, France). Then 329 individuals were sequenced in 150 paired-end reads on a Novaseq X Illumina sequencer (Novogene, Cambridge, UK). For *R. chacei* and *R. hybisae*, ten samples were included as triplicates in pooled libraries to enable estimation of the genotyping error rate. For *R. exoculata*, eight individuals were triplicated. Each plate comprised several negative controls to ensure the absence of contamination by foreign DNA.

### 2.3. De novo assembly, SNP calling and filtering

The *FastQC* v.0.11.9 software (Andrews, 2010) was used to assess the quality of the reads and the *process_radtags* module of *Stacks* v.2.52 (Catchen et al., 2011; Rochette et al., 2019) was run to demultiplex the individual files. Options *-c*, *-q* and *-r* were activated. These options (1) remove any read with an uncalled base, (2) discard reads with low quality scores and (3) rescue barcodes and RAD-Tags. The removal of adapter sequences with up to 2 mismatches was also performed with this program. As there is no reference genome available for *Rimicaris* species, two *de novo* assemblies were performed: one for *R. exoculata* and one for *R. chacei / R. hybisae*. Indeed, given the close genetic relationship between *R. chacei* and *R. hybisae* (Methou, Guéganton, et al., 2024; Teixeira et al., 2013; Vereshchaka et al., 2015) and in order to test their joint demographic history, they were processed together. The *r0.80* optimisation procedure detailed in Paris et al. (2017) and (Rochette & Catchen, 2017) (see Supplementary material for details) was applied to determine optimal *Stacks-M* and *-n* parameters values. The selected values were *-M* = *-n* = 4 and *-M* = *-n* = 6 for *R. exoculata* and *R. chacei / R. hybisae*, respectively (see Supplementary Figures S1 and S2). All the modules of the *Stacks* pipeline were then run sequentially on the entire dataset (i.e., including all sequenced individuals). However, the *cstacks* module was not run. Instead, *sstacks* with the-*-catalog* option was used to agglomerate all individuals on the loci catalogue obtained from the sub-sample of individuals (24 for R. exoculata and 40 for *R. chacei/R. hybisae*) used for parameter optimisation. For both assemblies, running the *populations* module with *-r* = 0.80 was a good compromise between the number of retrieved SNPs and missing data and thus retained. The maximum observed heterozygosity *(--max-obs-het*) to avoid paralog assembly and the minimum alternative allele count (*--min-mac*) required to keep a SNP at a locus were set to 0.80 and 10, respectively.

The following filtering steps were performed on the VCF file of genotypes obtained with *Stacks* v.2.52 using *VCFtools* (v.0.1.16) (Danecek et al., 2011) and *R* v.4.1.0 (R Core Team 2021) to eliminate SNPs linked to potential unfiltered paralogues, SNPs with too low coverage, SNPs erroneously genotyped between triplicated samples and SNPs and individuals with more than 10% of missing data (see Supplementary Material for details). Only the first SNP of each locus was retained for subsequent analyses to avoid linkage disequilibrium over short distance between SNPs of the same radtag. Outlier SNPs were filtered out using the *R* package *pcadapt* v. 4.3.3 (Luu et al., 2017; Privé et al., 2020), keeping the first two principal components of the analysis for each assembly (details in Supplementary material and Figures S3 and S4). All SNPs showing *p-values* < 0.05 after adjustment using *q-values* (Benjamini & Hochberg, 1995) were removed. The final filtered VCFs, which included 2020 independent SNPs for *R. chacei/R. hybisae* (mean coverage 69.89X) and 1258 independent SNPs for *R. exoculata* (mean coverage 69.32X), were produced using VCFtools individuals and SNPs white-lists. All Stacks assemblies were run on the bioinformatic cluster of the ABIMS platform hosted at the Station Biologique de Roscoff.

### 2.4. Population genetics analyses

To investigate the genetic structure of either *R. exoculata* or *R.chacei / R. hybisae* populations, we first performed a Principal Component Analysis (PCA) and a Discriminant Analysis of Principal Components (DAPC) for both assemblies using the *R* package *adegenet* v.2.1.3 (Jombart, 2008; Jombart & Ahmed, 2011). The DAPC enables the identification of the optimal number of genetic clusters (K) using a K-means algorithm (*find.clusters*) and the assignment of individuals to clusters (*dapc*). The function *find.clusters* was run for K ranging from 1 to 10, with 1000 iterations and 100 different starting points. The optimal number of clusters was identified as the one that minimised the BIC values. A cross-validation procedure was then used to identify the optimal number of principal components to retain to perform the DAPC. Coancestry analyses were performed using the maximum likelihood approach implemented in ADMIXTURE v.1.3 (Alexander & Lange, 2011; H. Zhou et al., 2011) for which datasets were formatted using PLINK v.1.9 (Chang et al., 2015; Purcell et al., 2007). ADMIXTURE was run 10 times, for K values ranging from 1 to 10, with the default parameters (admixture allowed). Tenfold cross-validation (CV) was applied to identify the number of clusters that minimised the average CV error value. Independent runs were then combined for the optimal K value using CLUMPP (Jakobsson & Rosenberg, 2007) as implemented in CLUMPAK (Kopelman et al., 2015). Finally, the *StAMPP R* package (Pembleton et al., 2013) was used to calculate pairwise fixation indices (F_st_) as estimated with theta proposed by (Weir & Cockerham, 1984). Fst_s_ were tested against zero by resampling loci (n = 1000 bootstraps) to obtain confidence intervals. Genetic diversity indices (observed and expected heterozygosities, nucleotide diversity π) and heterozygote deviations from Hardy-Weinberg equilibrium (F_is_) were calculated with the *populations* module of *Stacks* using all sites from all ddRAD tags of the final dataset. The direction and strength of migrations between *R.chacei/R. hybisae* populations were investigated using networks produced by DivMigrate (Sundqvist et al., 2016), with both the statistics *Gst* and Jost’s *D* calculated between pairs of populations under the null hypothesis of an N-islands model without demographic changes. The divMigrate-online pipeline (https://popgen.shinyapps.io/divMigrate-online/) was used for the analyses, with a Genepop file of 2,220 SNPs used as input. In the migration network, the number of migrants (*Nm*) was recalibrated between 0 and 1; 0 indicating no migrants per generation and 1 indicating a minimum of one migrant per generation exchanged from population A to population B. Migration asymmetry was then tested by resampling genotypes between pairs of populations with 1000 bootstraps.

### 2.5. Demographic history

We used the DILS software (Fraïsse et al., 2021) to make demographic inferences taking into account linked selection and heterogeneous migration across the genome. This was done on data from the two geographic populations of *R. chacei* (sampled along the MAR) and *R. hybisae* (sampled along the MCSC). The software employs an Approximate Bayesian Computation (ABC) framework to compare observed statistics, such as the number of bi-allelic sites shared (Ss) and fixed (Sf) between populations, the number of bi-allelic polymorphic sites within each species (Sx_A, Sx_B), nucleotide diversities (π or θ), Tajima’s D, F_ST_, D_xy_ and Pearson’s cross-correlations, and additional summary statistics from the joint site-frequency spectrum (jSFS), and then compares these with statistics simulated by isolation-population models, with and without migration (28 models). The software first identifies the best demographic model (from 4 main models: strict isolation (SI), ancient migration (AM), isolation-with-migration (IM) and secondary contact (SC)) and then identifies the best genomic model to test whether migration (2m) and effective size (2N) are heterogeneous along the genome. Finally, when necessary, it identifies the loci associated with barriers to gene flow. DILS also provides a goodness-of-fit statistic for all statistics used in the ABC analysis, determining how well the selected demographic model fits the genomic dataset. To run the two-population analysis, we used the DILS Snakemake pipeline (with Snakefile_2pop) on the ABIMS cluster. This requires the use of Python, R and PyPy, as specified in the DILS_2pop bash file together with a YAML file containing all the information on the biological parameters (e.g. species names, filtering preferences, mutation and recombination rates, prior distributions of population sizes, population splitting times and migration rates), and a cluster_2pop.json file with all the information on the module runs (e.g. allocated memory, run time). Before running DILS, the sequence data were filtered according to the maximum tolerated N, the minimum sequence length (L_min_) and the minimum number of individuals to be sampled at random from the sequence data for each investigated locus (N_min_). Statistics were then estimated from randomly selected subsamples of 1,000 loci with 1,000 replicates. Depending on the selected best population model, the estimated population parameters are as follows: Time since populations separated (Tsplit, all models); time since migration stopped (T_AM_, AM only); time since secondary contact and migration (T_SC_, SC only); population sizes of the ancestral population (N_A_) and sister populations (N_1_ and N_2_); whether these populations are constant, expanding or contracting (N_founder1_ and N_founder2_); and finally, the directional migration rates M_12_ and M_21_ estimated in the forward sense.

## 3. Results

### 3.1. Population genetic structure of Rimicaris exoculata along the northern MAR

No spatial genetic structure was detectable on the PCA for *Rimicaris exoculata* (Figure 2), but 2 individuals had outlier positions on the two first PCA axes (7.3% of the explained variance) (Figure 2A). Individuals from all vent fields overlapped with and without these outliers (Figure 2B). The DAPC procedure accordingly supported K = 1 as the optimal number of clusters (see: Supplementary Figure S5). The ADMIXTURE analysis also supported the hypothesis that *R. exoculata* populations form a single panmictic genetic group (see: Supplementary Figure S6 for the minimal average CV error value, and S7 for membership proportions from K = 2 to K = 5). At the population level, genetic diversity indices were similar across all sampled populations (see: Supplementary Table S2), and pairwise Fst values were in the order of 10^-3^ and did not significantly differ from zero, except for the pair Snake Pit/TAG, despite them being the closest geographically (Table 2).

**Figure 2:**
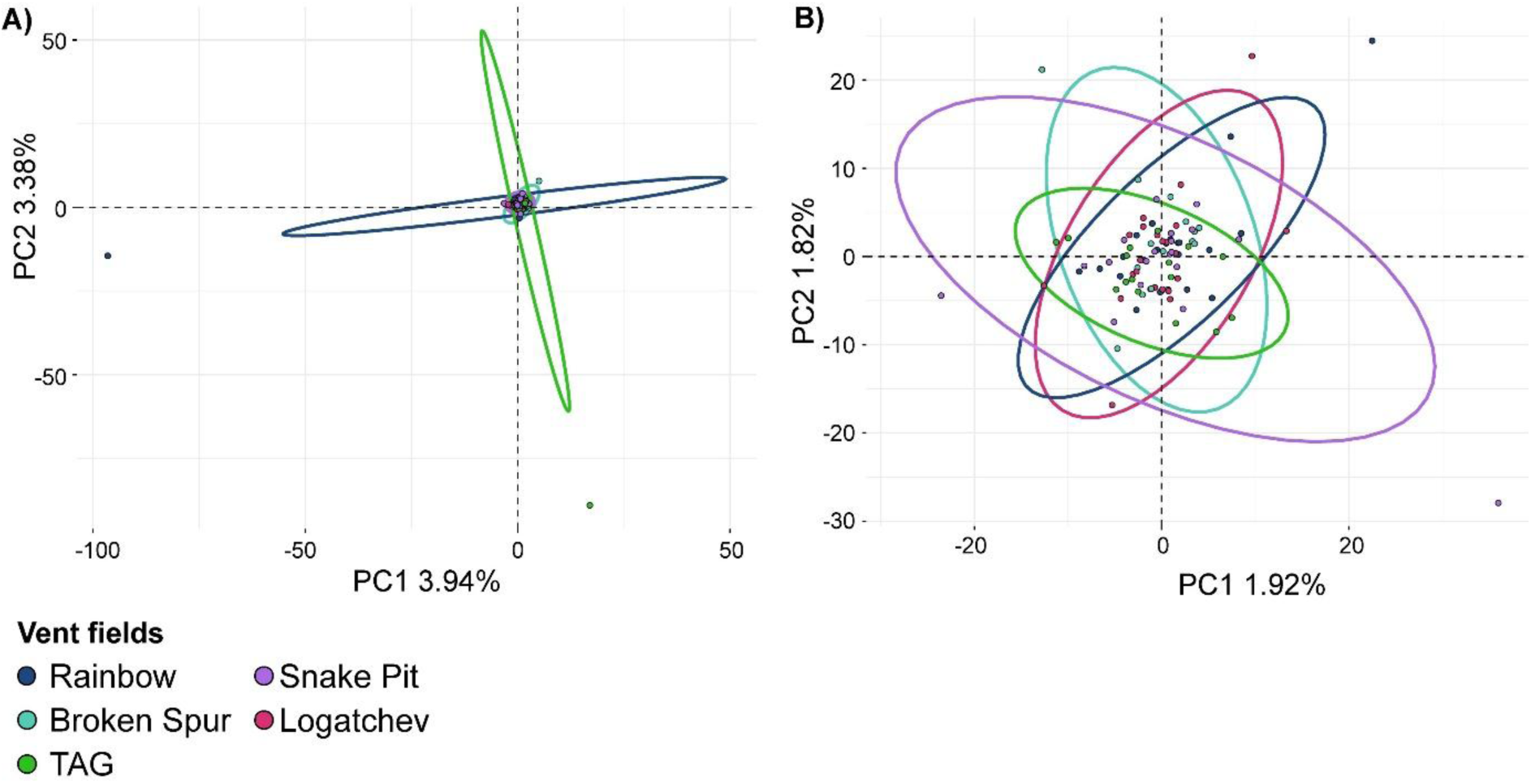
Population genetic structure of *Rimicaris exoculata* along the northern Mid-Atlantic Ridge **A.** Principal component analysis plot of genetic clusters considering all individuals **B.** or excluding 2 individuals that appear as outliers.

**Table 2:** Pairwise Fst values (below the diagonal with mean values and range under bracket) and Bonferroni corrected associated p-values (above the diagonal) between vent fields sampled for *R. exoculata* and *R. chacei / R. hybisae* assemblies. Significantly different from zero values are bold.

| *R. exoculata* |  |  |  |  |  |  |
| --- | --- | --- | --- | --- | --- | --- |
|  | Rainbow | Broken Spur | Logatchev | TAG | Snake Pit |  |
| Rainbow | - | 1.00 | 1.00 | 0.09 | 1.00 |  |
| Broken Spur | 0.0005 [-0.0026-0.0035] | - | 1.00 | 0.27 | 0.62 |  |
| Logatchev | -0.002 [-0.0042-0.0005] | -0.0013 [-0.0046-0.0021] | - | 1.00 | 1.00 |  |
| TAG | 0.0043 [0.0006-0.0079] | 0.0044 [-0.0001-0.0087] | 0.0023 [-0.0013-0.0059] | - | **0.00** |  |
| Snake Pit | 0.0005 [-0.0026-0.0037] | 0.0030 [-0.0006-0.0066] | -0.0001 [-0.0028-0.0028] | **0.0061 [0.0017-0.0107]** | - |  |
| *R. chacei* / *R. hybisae* |  |  |  |  |  |  |
|  | BrokenSpur | Lucky Strike | Snake Pit | TAG | BeeBe | Von Damm |
| BrokenSpur | - | 1.00 | 0.15 | 0.08 | **0.00** | **0.00** |
| Lucky Strike | 0.0003 [-0.0027-0.0033] | - | 1.00 | 1.00 | **0.00** | **0.00** |
| Snake Pit | 0.0026 [0.0004-0.0049] | 0.0007 [-0.002-0.0035] | - | 0.23 | **0.00** | **0.00** |
| TAG | 0.0029 [0.0006-0.0051] | 0.0016 [-0.0006-0.004] | 0.0015 [0.0002-0.003] | - | **0.00** | **0.00** |
| BeeBe | **0.0234 [0.0126-0.037]** | **0.0223 [0.0127-0.0353]** | **0.0231 [0.0133-0.0356]** | **0.0233 [0.0145-0.0351]** | - | 1.00 |
| Von Damm | **0.0246 [0.0144-0.0371]** | **0.0218 [0.0126-0.0336]** | **0.0243 [0.0148-0.0358]** | **0.0233 [0.0147-0.0336]** | -0.0005 [-0.0021-0.0013] | - |

### 3.2 Population genetic structure of the Rimicaris chacei / R. hybisae complex

Analysing the genetic structure of *R. chacei / R. hybisae* using PCA revealed a slight separation of individuals sampled along the MCSC (*R. hybisae*) and individuals sampled along the MAR (*R. chacei*) on the second axis, with some overlap. However, the first two principal components only explained a small proportion of genetic variance (2.42%; see Figure 3A). In the DAPC procedure, the BIC was minimised for K = 1, although K = 2 received some support (see: Supplementary Figure S5). Running the DAPC with K=2 on the *R. chacei/R. hybisae* genetic dataset using one discriminant function and 20 PC axes (cross-validation procedure) enabled about 24% of the genetic variance to be extracted, separating individuals into two groups: one comprising *R. chacei* individuals from all MAR vent sites and, the other containing *R. hybisae* individuals sampled in the Mid Cayman Spreading Centre (Figure 3B). Individual posterior membership probabilities in their respective clusters were all greater than 0.99, suggesting a weak but reliable population differentiation.

**Figure 3:**
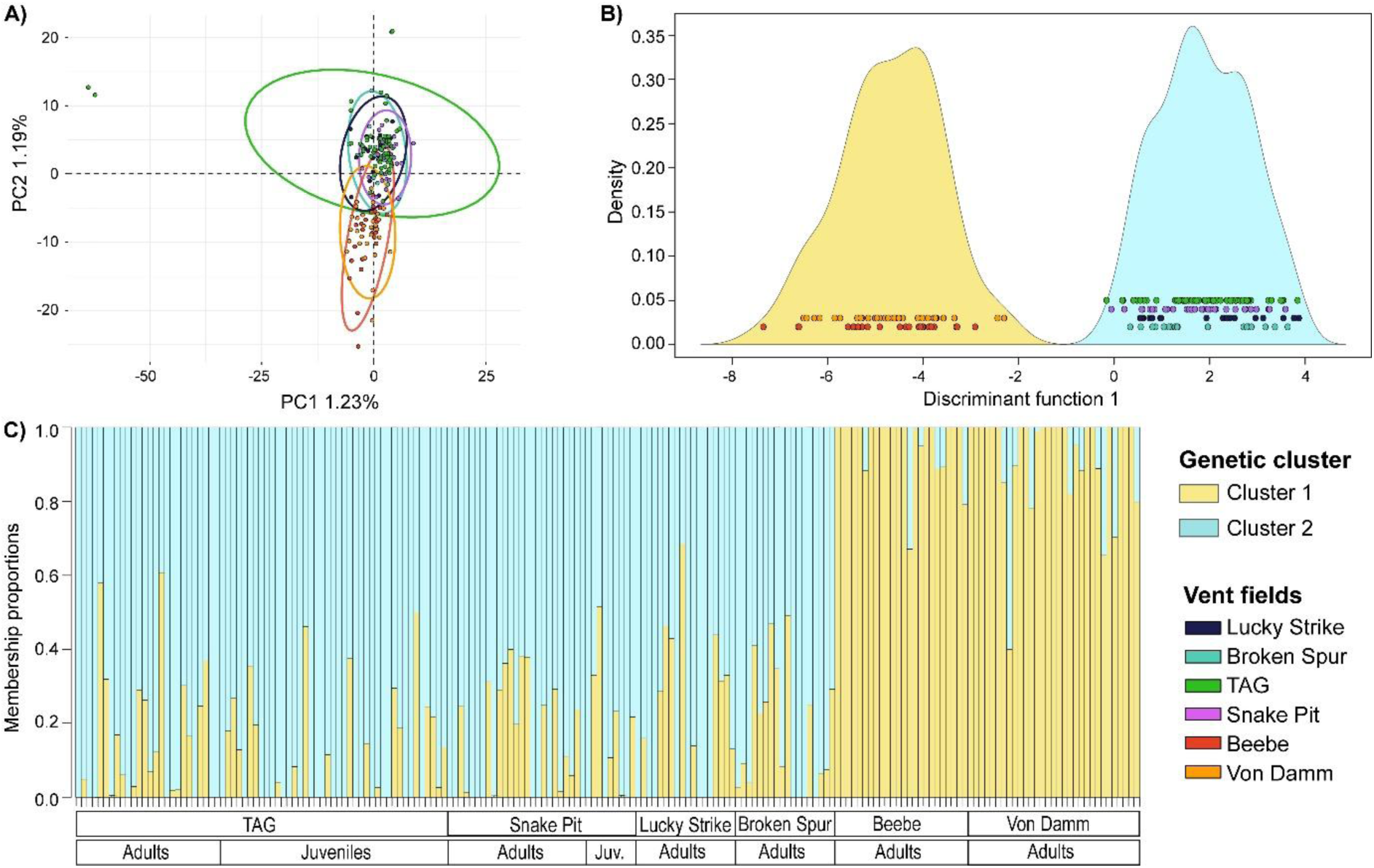
Population genetic structure of the *R. chacei* (Mid-Atlantic Ridge) and *R. hybisae* (Mid Cayman Spreading Center) complex of species. **A.** Principal component analysis plot of genetic clusters **A.** Discriminant Analysis of Principal Components (DAPC) genetic clusters **C.** ADMIXTURE membership proportions for *R. chacei/R. hybisae* complex of species.

Given the DAPC results with K = 2, we examined the ancestry coefficients in the context of two genetic clusters even though the K = 1 solution was preferred. Overall, *R. hybisae* individuals sampled at the MCSC form one group with ancestry memberships ranging from 60 to 100% for the first genetic cluster. Individuals of *R. chacei* sampled along the MAR belong to a second group with ancestry membership proportions ranging from 50 to 100% for the other genetic cluster. Several individuals had intermediate membership proportions, but allele frequency differences between the two genetic backgrounds were weak (very few outliers: only 10 above Fst=0.25) suggesting whetevthat these admixed individuals were not true hybrids (Figure 3C). Interestingly, one *R. hybisae* individual (Von Damm of MCSC) and three *R. chacei* individuals (TAG and Lucky Strike of MAR) were better assigned to the genetic cluster dominating the other geographic locality. More admixed individuals with the highest membership proportions for the alternative genetic cluster seemed to occur along the MAR than along the MCSC. Indeed, the proportion of non-admixed individuals was 0.70 for the MCSC shrimps and only 0.42 for the MAR shrimps.

Pairwise Fst values involving pairs between MCSC and MAR vent fields were around 0.02 and significantly different from zero, as also evidenced by the confidence intervals not including zero (Table 2). Genetic differentiation was, on the opposite, negligible and, at least one order of magnitude lower between either the MCSC sites or between the MAR sites. The divMigrate analysis estimated using the Gst statistic found a maximum gene flow between TAG and Snake Pit, indicating a strong connection between these two sites (Figure 4A). Using this statistic, a substantial gene flow with no asymmetry was also observed between the MCSC and MAR sites albeit less important than the one within each geographical region. However, the number of migrants estimated by Jost’s *D* suggested a much greater role for the MCSC Beebe site in supplying some of the MAR sites (Broken Spur and Lucky Strike) with migrants, indicating significant asymmetry in migration (Figure 4B). Examining more carefully allele frequency differences across 2020 loci revealed only one nearly fixed SNP between the *hybisae* and *chacei* populations and just 17 outlier SNPs with a frequency difference ranging from 0.25 to 0.5.

**Figure 4.**
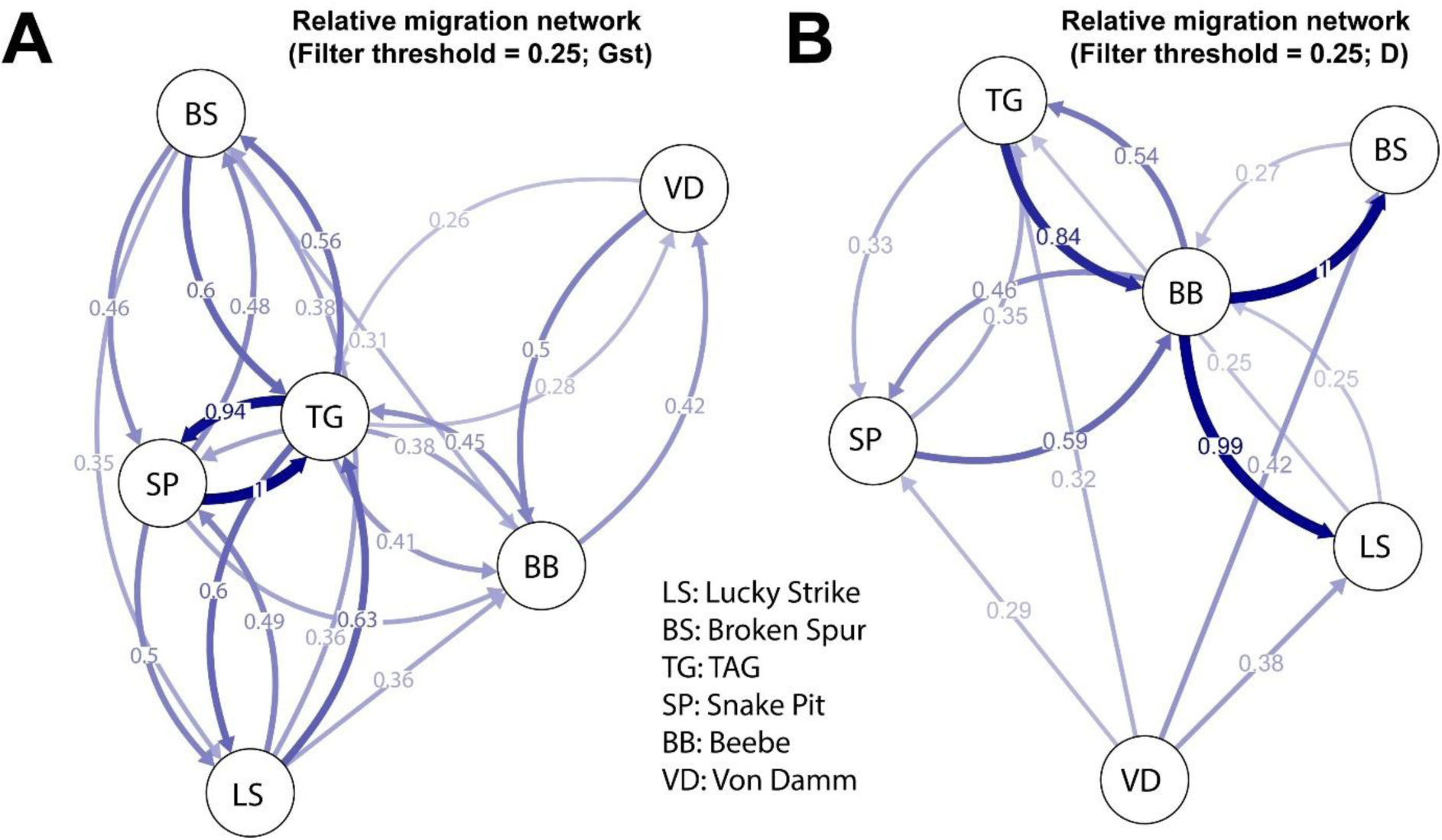
Relative migration networks of *R. chacei/hybisae* populations using divMigrate and based on **A.** Gst and **B.** Jost’s D statistics. Each node represents a population from a given vent field.

### 3.3. Demographic inferences for R. chacei and R. hybisae

As population genetics analyses revealed some genetic differentiation between the MAR *R. chacei* and MSCS *R. hybisae* shrimp populations, we investigated potential demographic scenarios connecting the two geographical groups using isolation-with-migration models (Figure 5). DILS results were found to be highly sensitive to the filtering conditions of the sequence dataset due to a trade-off between L_min_ and max_tolerated_N because of the very small observed divergence between the two taxa. The highest number of significant goodness-of-fit values for each population parameter of the best demographic model were found with a max_tolerated_N = 0.05 (no more than two gaps per locus in the sequence alignment) and Lmin = 140, resulting in 5,656 loci being examined in total. Decreasing N to zero greatly limited the number of loci used, whereas increasing it biases the estimation of genetic statistics (see Supplementary Table S3 for the number of loci retained and the p-values of goodness-of-fit for statistics estimated from different filtering attempts to select the best model). Due to the small number of loci used (N = 5,656), the number of sampled alleles per population and per locus was restricted to 20 in order to create 20 × 20 allele jSFS matrices that minimised the number of empty cases. In parallel, we increased the number of alleles used to build a 70 x 70 matrix to better estimate observed diversity statistics, which were subsequently used as an input in the ABC process. Consequently, for each filtering condition, we performed two DILS runs: one with jSFS and 20 alleles per population and locus, and one without jSFS and with 70 alleles per population and locus. This was done to check whether they produced similar genetic statistics and selected the same population model. The best population model selected for the analysis of the two *R. chacei* and *R. hybisae* populations was the SC_1m_2N (p-value migration vs isolation = 0.58, p-value SC vs IM = 0.63, p-value M_homo vs M_hetero = 0.64 and p-value N_hetero vs N_homo = 0.56: best model in bold) with a good congruence of statistics and goodness-of-fit of these statistics (Gof2) between the two DILS runs (with and without jSFS) (Table S4).

**Figure 5.**
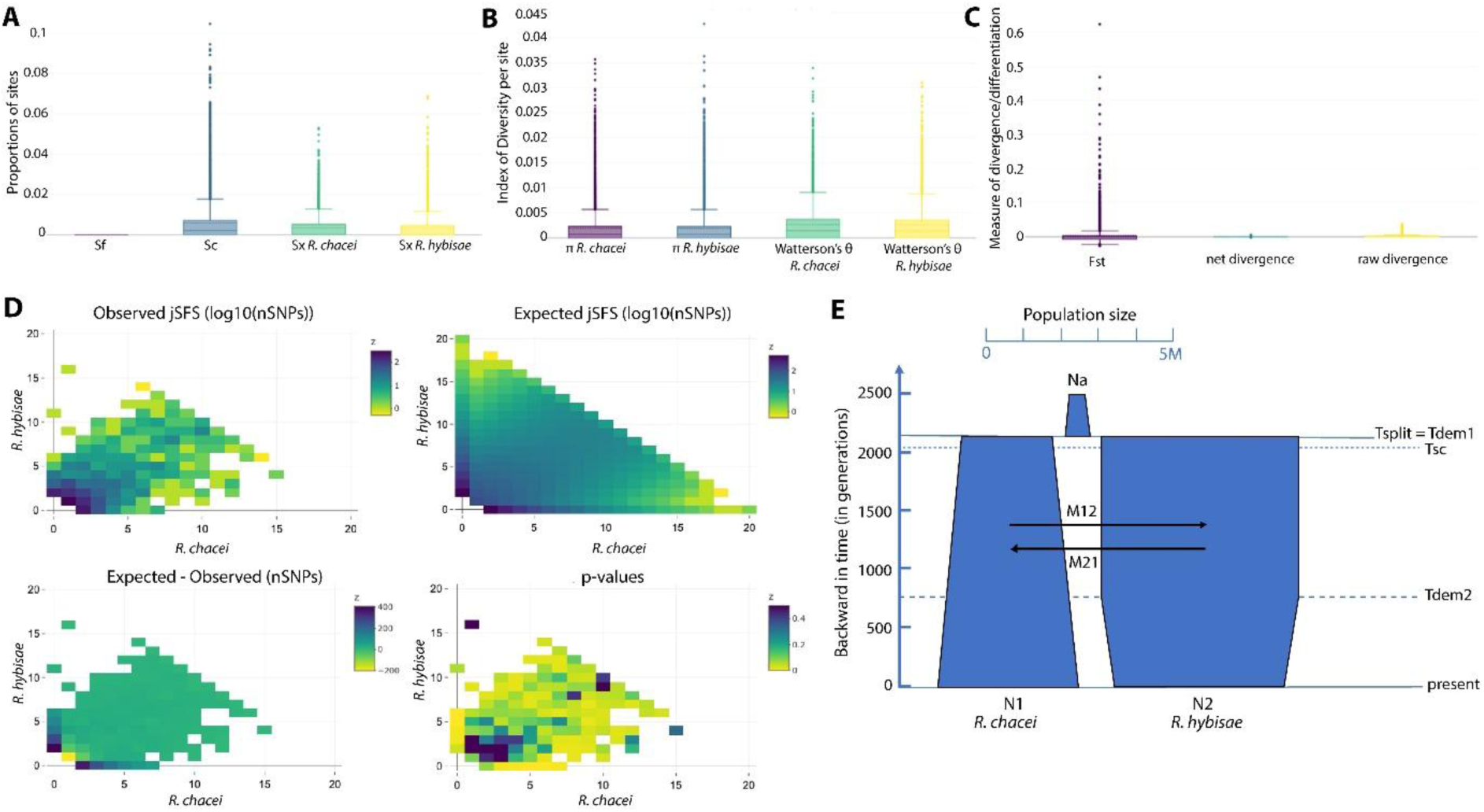
DILS analysis of the R. chacei/R. hybisae complex of species. **A.** Statistics displaying the number of bi-allelic sites shared (Ss) and fixed (Sf) between populations and the number of bi-allelic polymorphic sites (Sx) within each species. **B.** Nucleotide diversities (π or θ) **C.** Fst statistics **D.** Heatmap of both observed and expected 20×20 alleles joint site-frequency spectrum (jSFS) of the two shrimp populations (*R. chacei* vs *R. hybisae*) with residues and p-values of the model fit. **E.** Schematic graph of the evolution of the two ecotypes populations since their initial split following a SC_1m_2N model derived from the 2_populations DILS analysis. N_a_= Ancestral population size, N_1_ and N_2_: sister population sizes, T_split_= Time since population splitting, T_dem1_ and T_dem2_: Times since population expansion or contraction, T_sc_= Time since secondary contact, and M_12_ and M_21_: numbers of migrants exchanged per generation between populations 1 and 2 in forward time.

Statistics obtained from DILS on the whole filtered SNP data such as the number of fixed sites (*Sf*), the number of shared polymorphic sites (*Ss*) and, the number of species-specific polymorphic sites (*Sx_chacei, Sx_hybisae*) clearly supported the very close relationship of the two taxa (*R. chacei* and *R. hybisae*) with almost the same genetic diversity signatures and a very low divergence (Figure 5A). For both runs, the observed statistics also indicated that the two ecotype populations (*R. chacei* vs. *R. hybisae*) were close to panmixia, showing no genetic divergence (Da= 0.00003), much less population differentiation (Fst= 0.0039) than previously estimated and nearly identical low levels of genetic diversities (θ_N1_=0.0027, θ_N2_=0.0026), consistent with neutral evolution (Tajima’s D= −0.61 (N1) and - 0.56 (N2) (Figure 5B & C). However, it is worth noting that, unlike divergence estimates, the *Fst* distribution contains several outlier loci (Figure 5 C). Results first indicated that the two ecotype populations emerged from a very small ancestral population (0.018 million) and then increased rapidly in size (*N_1_*=3.7 million and *N_2_*=4.5 million), with the *R. hybisae* population initially being nearly twice the size of the *R. chacei* population (Figure 5D). Since the population split, the *R. chacei* population has steadily grown, while the *R. hybisae* population began contracting around 750 generations ago. The ecotype populations diverged very recently (*T_split_* around 2,100 generations ago) and they reunited almost instantly (*T_sc_* = 2,000 generations ago), suggesting that the allelic lineage sorting between the two ecotypes never truly began (Figure 5D). Migration rates between the two sister populations are high in both directions, but are slightly greater from *R. hybisae* to *R. chacei* (*M_12_* =21 vs *M_21_* =32 migrants per generation). As expected, given the lack of divergence, migration is homogeneous across the genome, with no barrier loci; however linked selection appears to be quite localised (β distribution of *Ne*: α=0.6, β=3.7 suggesting that the low recombination rate is restricted to a few linked loci) across the genome of the two species. The second-best model, selected by DILS with slightly fewer loci (*N* = 5,237), was AM_1m_2N. The population parameter means were almost identical to those of the first model for *N_A_*, *N_1_*, *N_2_*, *T_split_, M_12_*, *M_21_*, and *Shape_N_α* and *Shape_N_β*, but *T_dem_* and *T_AM_* (time at which migration ceased) coincided with *T_split_*. This indicates a very recent period of strict isolation, followed by population expansion, during which there was insufficient time for sorting alleles.

## 4. Discussion

### 4.1. Panmictic vent shrimp populations along the MAR

Genome-wide approaches employed here failed to reveal any genetic structure among the populations of *Rimicaris exoculata* or *R. chacei*, which are distributed over distances of 3,000 km and 2,000 km, respectively, along the Mid-Atlantic Ridge. Our results therefore confirm the findings of previous studies that used both mitochondrial COI gene and nuclear microsatellite markers (Teixeira et al., 2011, 2012, 2013), suggesting that populations of the two shrimp species are capable of exchanging larvae over long distances along the MAR. With the exception of a few outlier individuals (which may be due to either DNA contamination or the presence of immigrants from elsewhere), our genomic analysis, which was based on 1,258 (*R. exoculata*) and 2,220 (*R. chacei*) independent SNPs clearly showed complete mixing of individuals, regardless of their geographic origin. Consequently, the complete lack of genetic structure for both species along the Mid-Atlantic Ridge can only be explained by highly efficient migration over the ridge axis.

Unfortunately, the path taken by dispersing larval stages remains enigmatic, as the only evidence of their life outside the vent habitat is the photosynthetic signature of their tissue just after settlement (Pond et al., 2000). Their planktonic zoea larvae exhibit morphological characteristics indicative of an elongated larval phase (Hernández-Ávila et al., 2015) during which they develop in water masses, feeding on photosynthetic material originating from the surface production in order to accumulate lipid reserves (Methou et al., 2020; Pond et al., 2000). Early juveniles have been caught a few hundred meters above active vents along the ridge axis, suggesting that they actively migrate in the bathypelagic environment before recruitment (Herring & Dixon, 1998). They then return to vent sites as much larger juveniles than the larvae initially released into the water column (Methou et al., 2022). Whether larvae reach the photic zone of the ocean to feed on phytoplankton or feed at bathyal depths from sunken materials remains unknown. Vertical migration from hydrothermal vents to the surface has been suggested in the phenacolepadid limpet *Shinkailepas myojinensis* in the Western Pacific from in vivo larval experimentations (Yahagi et al., 2017) and was also demonstrated in the deep-sea seep mussel *Gigantidas childressi* (Arellano et al., 2014). Surface migration could, therefore, be the mechanism by which some vent species can achieve high dispersal along the ridge taking advantage of the stronger surface currents. Such mechanism was indeed suggested for the MAR vent gastropod *Divia briandi* (a phenacolepadid limpet like *S. myojinensis*) to explain its lack of genetic structure at the ridge scale, as determined by a mitochondrial gene (Yahagi et al., 2019). However, experimental work with the embryonic or larval stages of alvinocaridid shrimps suggests that the characteristics of these life stages may not be optimal to cope with the temperature and pressure of the photic ocean layer and that they may instead remain below the thermocline (Methou, Guéganton, et al., 2024; Tyler & Dixon, 2000).

The lack of genetic structure in vent shrimps contrasts sharply with recent population genomics studies that have been done on either the Atlantic vent mussels (Breusing et al., 2016; Van Der Heijden et al., 2012) or the vent gastropods *Lepetodrilus atlanticus* and *Peltospira smaragdina* (Portanier et al. in press), which all display highly structured populations along the Mid-Atlantic Ridge. Although the two gastropod genera have lecithotrophic larvae with likely deep-water dispersal (Chen et al., 2025; Mullineaux et al., 1996), *Bathymodiolus* vent mussels possess planktotrophic larvae with a relatively long pelagic larval duration. The difference between shrimps and mussels remains unclear, this could be due to the shrimp larvae being better at selecting the water layer to disperse in, or the result of environmental factors limiting the mussels’ dispersal, or the mussels having a shorter pelagic larval duration than shrimps.

Vent shrimp populations studied here all appear to contribute migrants that could reach other sites and participate in the renewal of other local populations, apparently without suffering regional connectivity restrictions, as observed in other vent species (Gollner et al., 2016; Methou, Ogawa, et al., 2024; Yahagi et al., 2019; Y. Zhou et al., 2022). Connectivity networks may however be affected by the impacts of mining activities, which are not limited to the seafloor, but potentially far reaching in the water column (Menini & Van Dover, 2019). Indeed, the persisting lack of knowledge on shrimp planktonic larval stages regarding their location above the seafloor, their specific physiological requirements, or their reliance on specific navigation cues to re-establish themselves at vents, make predictions on the impacts of mining extractive activities on connectivity networks highly uncertain.

### 4.2. Caribbean-Atlantic connectivity of the Rimicaris chacei / R. hybisae complex

The MCSC constitutes a unique and isolated vent biogeographic province in the Atlantic. It has one of the lowest faunal richness and comprises species that are unique to the region but also species that are shared with methane seeps in the Gulf of Mexico (Georgieva et al., 2023; Linse et al., 2019; Plouviez et al., 2015). Our results revealed a strong connection between the MCSC and the MAR vent fauna: the dominant MCSC taxon, *R. hybisae*, shares a population history with the MAR *R. chacei* shrimp population, exhibiting minimal genetic differentiation and contemporary gene flow. In addition, the lack of a fixed divergence both at the mitochondrial (Methou, Guéganton, et al., 2024) and nuclear (present study) genome scales further supports the hypothesis of a single species cross-exchanging genes between the MAR and MCSC. Based on these results, it is now clear that *R. chacei* and *R. hybisae* represent two distinct phenotypes or forms of a single species and we here synonymise them. *Rimicaris chacei* is the earlier name and has precedence over *R. hybisae* (see: section 5. Systematics).

The demogenetic analysis conducted by DILS (Fraïsse et al., 2021) clearly indicated the presence of a single shrimp population, with the *hybisae* and *chacei* forms exhibiting identical genetic diversity and minimal population differentiation. This contrasted with the geographic variation detected by the discriminating analysis of DAPC on the whole set of SNPs obtained from the shrimp genome. As DILS employed subsamples of 1,000 loci at each step of the ABC analysis, some population divergence was probably not detected – most likely because differentiated SNPs were rare – which made it easy to reject the strict isolation model but difficult to select the appropriate isolation-with-migration model. While genetic variables best fit the secondary contact model (SC_1m_2N), the DILS_2populations analysis makes it difficult to choose between models of secondary contact (SC), isolation with migration (IM) and ancient migration (AM) depending on the filtering parameters. Nevertheless, all models with migration agree that the two shrimp ecotypes have been separated for only a very short time, approximately less than 2,000 generations. This is much shorter than the estimated divergence time between geographic populations of several vent species including gastropods, mussels, polychaetes and barnacles, in back-arc basins of the southwestern Pacific (Tran Lu Y et al., 2022), or between southern and northern populations of several species in the East Pacific Rise (Matabos et al., 2011; Plouviez et al., 2009). It is therefore consistent with the hypotheses of ongoing migration between the MCSC and MAR, and of a rapid expansion following their initial separation, to explain the low genetic diversity observed despite large population sizes.

All isolation-with-migration models also indicated that populations of both *chacei* and *hybisae* forms increased rapidly by at least one order of magnitude following the split of the ancestral population. A possible explanation for this sudden and simultaneous population expansion could be the recent and shared acquisition of a trophic chemosynthetic symbiosis, which has been ecologically successful at several hydrothermal vent fields across the North Atlantic and Caribbean Sea (Methou et al., 2020; Methou, Guéganton, et al., 2024). Furthermore, *R. chacei* populations are growing faster possibly at the expense of *R. hybisae* populations, with reliable asymmetry in migration from the MCSC to the MAR, regardless of the chosen model. Although *R. chacei* populations are currently far smaller than those of their ecological challenger, *R. exoculata,* at MAR vents, this population increase is partly validated by the discovery of dense aggregations of *R. chacei* juveniles in nurseries at the TAG and Snake Pit vent fields despite a high post-settlement mortality (Hernández-Ávila et al., 2022; Methou et al., 2022). However, even if there is currently a massive recruitment of *R. chacei* juveniles at MAR vents, their genetic signatures are not different from those of adults and are not indicative of first- or second generation of migrants from MCSC. Consequently, the connection between the shrimp populations of the MCSC and MAR vents is probably indirect and should require intermediate active sites.

The bidirectionality of gene flow between the MAR and MCSC shrimp populations calls into question the lack of *R. exoculata* colonization at the MCSC vent fields. This can be viewed as either the migration of *R. chacei* forms being more asymmetrical than predicted by the model or, that *R. exoculata* is less able to migrate to the MCSC. Firstly, the co-occurring *Rimicaris* shrimps from the MAR exhibit different distributions and likely distinct demographic histories within the Atlantic Ocean, which may be related to the use of different dispersal routes. Despite sharing similar larval characteristics in the early stages (Hernández-Ávila et al., 2015), *R. exoculata* and *R. chacei* clearly differ in their juvenile sizes at settlement as well as in their isotopic niches that segregate in their carbon source in the earliest stages suggesting distinct larval histories (Methou et al., 2020). Secondly, co-occurring vent species often exhibit different connectivity patterns even between closely related species within the same genus due to specific combination of biological traits and habitat requirements (Chen et al., 2025; Methou et al., 2025; Poitrimol et al., 2022). In the case of the *Peltospira* species along the MAR, the restricted range of *P. gargantua* to the Hydra and Falkor EMARK vent fields, as opposed to the broader distribution of *P. smaragdina* across the northern MAR, has been attributed to specific environmental filters related to the geochemistry of these vents rather than to larval dispersal ability (Chen et al., 2025). Therefore, the two species likely have both different plastic responses to environmental variations and distinct larval life capabilities, resulting in different vertical distributions in the water column. This could prevent *R. exoculata* from reaching the Caribbean Sea during its dispersal phase.

### 4.3. Evolutionary implications of becoming a species with two distinct ecotypes

Contemporary exchanges of migrants between the MAR and MCSC shrimp populations suggests a strong ecological plasticity of the ‘panmictic’ species in terms of trophic behavior and habitat use. The *hybisae* form predominantly feeds on chemoautotrophic production, possibly from its cephalothoracic symbionts. It lives in dense aggregates and shows an enlarged cephalothorax. The *chacei* form shows limited anatomical changes during its metamorphosis with a less developed cephalothorax. It has a mixed diet, partially uses chemoautotrophic bacterial production, and lives in smaller groups (Methou et al., 2022; Methou, Guéganton, et al., 2024; Nye et al., 2013). It has been suggested that individuals of *R. chacei* could have a limited “critical period” during their metamorphosis that determines their developmental trajectory, whether an energetically viable symbiosis has been established before or after this window of competency (Methou, Guéganton, et al., 2024). Among factors that can affect their developmental trajectories, competition for space with *R. exoculata* on the MAR for access to suitable vent fluid sources and symbionts during the juvenile stages may drive development towards the *chacei* form. This is supported by the strong spatial segregation of the juvenile stages of *R. exoculata* and *R. chacei*; the latter is confined to more distant nursery habitats (Hernández-Ávila et al., 2022; Methou et al., 2022).

However, competition for a trophic niche should not entirely eliminate the *hybisae* form at MAR vent sites. Indeed, even in the absence of their *R. exoculata* competitor, *R. chacei* shrimps from the Lucky Strike vent field do not exhibit the inflated cephalothorax of the *hybisae* form. Thus, the slight genetic differences between the *hybisae* and *chacei* forms depicted by the DAPC analysis could be adaptive, regulating the molecular dialogue between the shrimp and its symbionts, and/or stabilising the morphological change of *R. chacei* along the MAR. It is worth noting that none of the sequenced *R. chacei* adults and juveniles had more than a 70% of ‘*hybisae’* ancestry.

Nevertheless, trophic plasticity also appears to be a common trait in *hybisae* populations on the MCSC. Individuals found on the periphery exhibit mixotrophic behaviour similar to that observed in *R. chacei*, feeding on the exoskeletons (moults) of their conspecifics (Versteegh et al., 2023). This duality of feeding behaviour (symbiotrophy vs. carnivory or scavenging: (Methou et al., 2020) could therefore be rooted in the plasticity and polymorphism of the species and, expressed differentially depending on habitat conditions and access to resources. Other cases of plasticity in relation to their habitat are already known in vent ecosystems, such as the “short-fat” or a “long-skinny” morphotypes of *Ridgeia piscesae* (Tunnicliffe et al., 2014) or the substrate-dependent shell morphologies of *Lepetodrilus nux* (Chen & Watanabe, 2020). Hence, some species possess distinct adult ecotypes able to colonise different habitats thanks to a complex interplay of genetic, epigenetic, and environmental determinants (Barrett & Schluter, 2008; Gore et al., 2018; Krishnan & Rohner, 2017; Schwander et al., 2010). The recent expansion of *R. chacei* can have led to allele surfing in the MAR populations that is known to promote rapid adaptation from the standing genetic variation (Barrett & Schluter, 2008; Excoffier & Ray, 2008), and hence the subtle genetic differentiation observed between its geographic populations. Our results highlight phenotypic and trophic plasticity as an important adaptive strategy to survive and thrive in hydrothermal vents, one of the Earth’s most dynamic ecosystems.

## 5. Systematics

**Family Alvinocarididae Christoffersen, 1986 Genus Rimicaris Williams & Rona, 1986**

**Type species:** *Rimicaris exoculata* Williams & Rona, 1986

*Rimicaris chacei* (Williams & Rona, 1986)

*Rimicaris hybisae* Nye *et al.,* 2012; **syn. nov.**

### Diagnosis

Adapted from (Komai & Segonzac, 2008) and (Nye et al., 2012). Carapace with broadly triangular or rounded antennal lobe; pterygostomial lobe broadly rounded, less produced. Scaphognathite of maxilla and caridean lobe of first maxilliped bearing numerous plumose seta-like structure on dorsal and ventral surface; exopodal flagellum on first maxilliped completely reduced. Appendix masculina tapering distally with 4-5 (*chacei* form) or 7–8 (*hybisae* form) spiniform setae restricted to tip. Uropodal protopod posterolateral projection triangular with acute tip.

<u>Characters specific to the *chacei* form on the MAR:</u> Rostrum bluntly triangular, overreaching anterior margins of eye-talks or tips of antennal lobes by less than half of its length; ventral surface slightly convex. Antennae suboperculate, directed downward; distolateral tooth subacute or blunt (Komai & Segonzac, 2008).

<u>Characters specific to the *hybisae* form on the MCSC:</u> Anterolateral region of the carapace inflated. Rostrum reduced to broadly rounded lobe, nearly reaching, reaching, or slightly overreaching anterior margins of medially-fused eyes; ventral surface flat or slightly convex. Four-lobed dorsal organ; lobes fused anteriorly; posterior lobes with paired <u>“</u>pores<u>”</u>. Antennae not operculiform; distolateral tooth subacute or blunt (Nye et al., 2012).

### Remarks

We here synonymise *R. hybisae* Nye *et al*., 2012 with *R. chacei* (Williams & Rona, 1986). *Rimicaris hybisae* was described as morphologically most similar to *R. exoculata*, *R. kairei* and *R. chacei* because of the reduction of the rostrum, the presence of plumose-seta like structures on the dorsal and ventral surfaces of enlarged mouthparts and an appendix masculina armed with distal setae only (Nye et al., 2012). However, it can be distinguished from *R. exoculata* and *R. kairei* by the slightly less inflated carapace and from *R. chacei* by the more inflated anterolateral region of the carapace, the possession of a four-lobed dorsal organ, and of an acute tip on the uropodal protopod (Nye et al., 2012). Recent work has shown that carapace inflation and the acquisition of a four-lobed dorsal organ are morphological characteristics acquired during metamorphosis with a differentiation of developmental trajectories between *R. chacei* and *R. hybisae* which only appears during this developmental phase with indistinguishable morphologies between the two species at juvenile stages (Methou, Guéganton, et al., 2024). Moreover, several morphological features of their internal anatomy, with equivalent stomach size and mouthparts surface areas at adult stage, support the conspecificity of *R. chacei* and *R. hybisae* (Guéganton, et al., 2024). Our work, using multiple single nucleotide positions (SNPs) markers support the existence of a single genetic cluster for *R. hybisae* and *R. chacei*, already suggested by previous work based on partial fragment of the mitochondrial COI gene (Teixeira et al., 2013; Vereshchaka et al., 2015), and also on partial fragments of 16S, 18S, 28S and H3 genes (Methou, Guéganton, et al., 2024).

### Distribution and habitat

The typical form is known at several hydrothermal vents on the MAR including Lucky Strike, Rainbow, Broken Spur, TAG, Hydra, Snake Pit, Puy des Folles, Logatchev and Ashadze vent fields, at depths of 1700–4088 m (Cambon, 2023; Komai & Segonzac, 2008). The form *hybisae* is known from the Von Damm and Beebe vent fields on the MCSC at depths of 2300–4960 m (Nye et al., 2012). Adults are observed both in dense aggregations of thousands of individuals around vent chimneys and scattered at the periphery on the MCSC (Nye et al., 2013) but mostly observed as scattered individuals or in small aggregations between rocks or behind mussel beds on the MAR (Methou et al., 2022). Juveniles were also observed in dense aggregations close to low diffuse emissions on the MAR (Methou et al., 2022). A relatively dense aggregation of *R. chacei* was also recently observed at the TAG vent field in November 2023 (Figure S8; Bicose 3 expedition; https://doi.org/10.17600/18002399).

## Supporting Information

Additional supporting information may be found in the online version of the article at the publisher’s website.

## Data Availability

The ddRAD sequences generated in this study are available from the European Nucleotide Archive database (project PRJEB61450).

## Acknowledgment

We thank the captain, crew, and scientists on-board R/V *Pourquoi Pas?* during research expeditions SERPENTINE (https://doi.org/10.17600/7030030), MOMARDREAM (https://doi.org/10.17600/7030060), HERMINE (https://doi.org/10.17600/17000200) and BICOSE2 (https://doi.org/10.17600/18000004), on-board R/V *Atalante* during expeditions TRANSECT (https://doi.org/10.17600/18000513), MOMARSAT20 (https://doi.org/10.17600/18000684), and MOMARSAT21 (https://doi.org/10.17600/18001296), on-board R/V *Yokosuka* during expeditions YK13-05, and on-board R/V *James Cook* during expedition JC82. We extend the same to the pilots and technical teams of ROVs *Victor 6000* and *Isis* as well as the HOVs *Nautile* and *Shinkai 6500* during those expeditions. We gratefully acknowledge the cruise PIs for leading the expeditions:); Yves Fouquet, Ifremer (SERPENTINE & HERMINE); Françoise Gaill, Sorbonne Universités and Javier Escartin, ENS (MOMARDREAM); Cécile Cathalot and Ewan Pelleter, Ifremer (HERMINE); Marie-Anne Cambon, Ifremer (BICOSE2); Nadine Le Bris, Sorbonne Universités (TRANSECT); Pierre-Marie Sarradin and Julien Legrand, Ifremer (MOMARSAT20); Marjolaine Matabos and Jozée Sarrazin, Ifremer (MOMARSAT21); Ken Takai, JAMSTEC (YK13-05) and Jon Copley, NOC (JC82). Permission for sampling R. hybisae specimens in territorial waters of the Cayman Islands was issued by the appropriate UK authorities to R/V Yokosuka during YK13-05. We thank Dr. Jon Copley (University of Southampton, UK) for providing *R. hybisae* samples from Von Damm collected during the JC82 expedition for our study, as well as Dr. Adrian Glover for the picture of *R. hybisae* (Figure 1C) took on board of the same expedition. This work benefited from access to the EMBRC-France and Biogenouest Genomer platform at the Station Biologique de Roscoff for ddRAD library size selection and quality control. We are also grateful to the Roscoff Bioinformatics platform ABiMS for access to the cluster for analyses and data storage. This project/work contributes the EMSO-Azores (https://www.emso-fr.org) regional node of the EMSO ERIC Research Infrastructure (https://emso.eu/) and was supported by the French Oceanographic Fleet (FOF https://www.flotteoceanographique.fr/en).

## References

Adams, D. K., Arellano, S. M., & Govenar, B. (2012). Larval dispersal: Vent life in the water column. Oceanography, 25(1), 256–268. 10.5670/oceanog.2012.24

Alexander, D. H., & Lange, K. (2011). Enhancements to the ADMIXTURE algorithm for individual ancestry estimation. BMC Bioinformatics, 12(1). 10.1186/1471-2105-12-246

Andrews, S. (2010). FastQC: a quality control tool for high throughput sequence data. Version 0.10, 1.

Arellano, S. M., Van Gaest, A. L., Johnson, S. B., Vrijenhoek, R. C., & Young, C. M. (2014). Larvae from deep-sea methane seeps disperse in surface waters. Proceedings of the Royal Society B: Biological Sciences, 281(1786), 20133276. 10.1098/rspb.2013.3276

Barrett, R. D. H., & Schluter, D. (2008). Adaptation from standing genetic variation. Trends in Ecology and Evolution, 23(1), 38–44. 10.1016/j.tree.2007.09.008

Beaulieu, S. E., Baker, E. T., & German, C. R. (2015). Where are the undiscovered hydrothermal vents on oceanic spreading ridges? Deep Sea Research Part II: Topical Studies in Oceanography, 121, 202–212. 10.1016/j.dsr2.2015.05.001

Benjamini, Y., & Hochberg, Y. (1995). Controlling the False Discovery Rate: A Practical and Powerful Approach to Multiple Testing. Journal of the Royal Statistical Society Series B: Statistical Methodology, 57(1), 289–300. 10.1111/j.2517-6161.1995.tb02031.x

Brelsford, A., Dufresnes, C., & Perrin, N. (2016). High-density sex-specific linkage maps of a European tree frog (Hyla arborea) identify the sex chromosome without information on offspring sex. Heredity, 116(2), 177–181. 10.1038/hdy.2015.83

Breusing, C., Biastoch, A., Drews, A., Metaxas, A., Jollivet, D., Vrijenhoek, R. C., Bayer, T., Melzner, F., Sayavedra, L., Petersen, J. M., Dubilier, N., Schilhabel, M. B., Rosenstiel, P., & Reusch, T. B. H. (2016). Biophysical and Population Genetic Models Predict the Presence of “Phantom” Stepping Stones Connecting Mid-Atlantic Ridge Vent Ecosystems. Current Biology, 26(17), 2257–2267. 10.1016/j.cub.2016.06.062

Breusing, C., Vrijenhoek, R. C., & Reusch, T. B. H. (2017). Widespread introgression in deep-sea hydrothermal vent mussels. BMC Evolutionary Biology, 17(1), 1–10. 10.1186/s12862-016-0862-2

Cambon, M.-A. (2023). Rapport de campagne BICOSE 3 2023. https://campagnes.flotteoceanographique.fr/campagnes/18002399/fr/

Cambon-Bonavita, M.-A., Aubé, J., Cueff-Gauchard, V., & Reveillaud, J. (2021). Niche partitioning in the Rimicaris exoculata holobiont: The case of the first symbiotic Zetaproteobacteria. Microbiome, 9(1), 1–16. 10.1186/s40168-021-01045-6

Catchen, J. M., Amores, A., Hohenlohe, P., Cresko, W., & Postlethwait, J. H. (2011). Stacks: Building and Genotyping Loci De Novo From Short-Read Sequences. G3 Genes|Genomes|Genetics, 1(3), 171–182. 10.1534/g3.111.000240

Chang, C. C., Chow, C. C., Tellier, L. C., Vattikuti, S., Purcell, S. M., & Lee, J. J. (2015). Second-generation PLINK: Rising to the challenge of larger and richer datasets. Gigascience, 4(1). 10.1186/s13742-015-0047-8

Chen, C., Pradillon, F., Diaz-Recio Lorenzo, C., & Alfaro-Lucas, J. M. (2025). Integrative taxonomy of two new peltospirid gastropods from Mid-Atlantic Ridge hot vents, including a potentially symbiotic species. Zoological Journal of the Linnean Society, 204(2), zlaf055. 10.1093/zoolinnean/zlaf055

Chen, C., & Watanabe, H. K. (2020). Substrate-dependent shell morphology in a deep-sea vetigastropod limpet. Marine Biodiversity, 50(6), 104. 10.1007/s12526-020-01135-y

Cuvelier, D., Matabos, M., & Sarrazin, J. (2025). Peltospira smaragdina gastropod assemblages as pioneers in Atlantic hydrothermal vent community succession. Marine Biology, 172(9), 136. 10.1007/s00227-025-04695-4

Daguin Thiebaut, C., Ruault, S., Roby, C., Broquet, T., Viard, F., & Brelsford, A. (2021). Construction of individual ddRAD libraries v1. Springer Science and Business Media LLC. 10.17504/protocols.io.bv4tn8wn

Danecek, P., Auton, A., Abecasis, G., Albers, C. A., Banks, E., DePristo, M. A., Handsaker, R. E., Lunter, G., Marth, G. T., Sherry, S. T., McVean, G., Durbin, R., & 1000 Genomes Project Analysis Group. (2011). The variant call format and VCFtools. Bioinformatics, 27(15), 2156–2158. 10.1093/bioinformatics/btr330

Desbruyères, D., Almeida, A., Biscoito, M., Comtet, T., Khripounoff, A., Le Bris, N., Sarradin, P. M., & Segonzac, M. (2000). A review of the distribution of hydrothermal vent communities along the northern Mid-Atlantic Ridge: Dispersal vs. environmental controls. In M. B. Jones, J. M. N. Azevedo, A. I. Neto, A. C. Costa, & A. M. F. Martins (Eds.), Island, Ocean and Deep-Sea Biology (pp. 201–216). Springer Netherlands. 10.1007/978-94-017-1982-7_19

Diehl, A., & Bach, W. (2020). MARHYS (MARine HYdrothermal Solutions) Database: A Global Compilation of Marine Hydrothermal Vent Fluid, End Member, and Seawater Compositions. Geochemistry, Geophysics, Geosystems, 21(12). 10.1029/2020gc009385

Faure, B., Jollivet, D., Tanguy, A., Bonhomme, F., & Bierne, N. (2009). Speciation in the deep sea: Multi-locus analysis of divergence and gene flow between two hybridizing species of hydrothermal vent mussels. PLoS ONE, 4(8), 1–15. 10.1371/journal.pone.0006485

Fraïsse, C., Popovic, I., Mazoyer, C., Spataro, B., Delmotte, S., Romiguier, J., Loire, É., Simon, A., Galtier, N., Duret, L., Bierne, N., Vekemans, X., & Roux, C. (2021). DILS: Demographic inferences with linked selection by using ABC. Molecular Ecology Resources, 21(8), 2629–2644. 10.1111/1755-0998.13323

Georgieva, M. N., Rimskaya-Korsakova, N. N., Krolenko, V. I., Van Dover, C. L., Amon, D. J., Copley, J. T., Plouviez, S., Ball, B., Wiklund, H., & Glover, A. G. (2023). A tale of two tubeworms: Taxonomy of vestimentiferans (Annelida: Siboglinidae) from the Mid-Cayman Spreading Centre. Invertebrate Systematics, 37(3), 167–191. 10.1071/is22047

Gollner, S., Stuckas, H., Kihara, T. C., Laurent, S., Kodami, S., & Arbizu, P. M. (2016). Mitochondrial DNA Analyses Indicate High Diversity, Expansive Population Growth and High Genetic Connectivity of Vent Copepods (Dirivultidae) across Different Oceans. PLoS ONE, 11(10), 1–23. 10.1371/journal.pone.0163776

Gore, A. V., Tomins, K. A., Iben, J., Ma, L., Castranova, D., Davis, A. E., Parkhurst, A., Jeffery, W. R., & Weinstein, B. M. (2018). An epigenetic mechanism for cavefish eye degeneration. Nature Ecology and Evolution, 2(7), 1155–1160. 10.1038/s41559-018-0569-4

Guéganton, M., Methou, P., Aubé, J., Noël, C., Rouxel, O., Cueff, V., Gayet, N., Durand, L., Pradillon, F., & Cambon-Bonavita, A. (2024). Symbiont Acquisition Strategies in Post-Settlement Stages of Two Co-Occurring Deep-Sea Rimicaris Shrimp. Ecology and Evolution.

Hernández-Ávila, I., Cambon-Bonavita, M. A., & Pradillon, F. (2015). Morphology of first zoeal stage of four genera of alvinocaridid shrimps from hydrothermal vents and cold seeps: Implications for ecology, larval biology and phylogeny. PLoS ONE, 10(12), 1–27. 10.1371/journal.pone.0144657

Hernández-Ávila, I., Cambon-Bonavita, M.-A., Sarrazin, J., & Pradillon, F. (2022). Population structure and reproduction of the alvinocaridid shrimp Rimicaris exoculata on the Mid-Atlantic Ridge: Variations between habitats and vent fields. Deep Sea Research Part I: Oceanographic Research Papers, 186(June), 3–5. 10.1016/j.dsr.2022.103827

Herring, P. J., & Dixon, D. R. (1998). Extensive deep-sea dispersal of postlarval shrimp from a hydrothermal vent. Deep-Sea Research Part I: Oceanographic Research Papers, 45(12), 2105–2118. 10.1016/S0967-0637(98)00050-8

Hurtado, L. A., Lutz, R. A., & Vrijenhoek, R. C. (2004). Distinct patterns of genetic differentiation among annelids of eastern Pacific hydrothermal vents. Molecular Ecology, 13(9), 2603–2615. 10.1111/j.1365-294X.2004.02287.x

Jakobsson, M., & Rosenberg, N. A. (2007). CLUMPP: A cluster matching and permutation program for dealing with label switching and multimodality in analysis of population structure. Bioinformatics, 23(14), 1801–1806. 10.1093/bioinformatics/btm233

Jombart, T. (2008). *adegenet*: A R package for the multivariate analysis of genetic markers. Bioinformatics, 24(11), 1403–1405. 10.1093/bioinformatics/btn129

Jombart, T., & Ahmed, I. (2011). *adegenet 1.3-1*: New tools for the analysis of genome-wide SNP data. Bioinformatics, 27(21), 3070–3071. 10.1093/bioinformatics/btr521

Komai, T., & Segonzac, M. (2008). Taxonomic Review of the Hydrothermal Vent Shrimp Genera Rimicaris Williams & Rona and Chorocaris Martin & Hessler (Crustacea: Decapoda: Caridea: Alvinocarididae). Journal of Shellfish Research, 27(1), 21–41. 10.2983/0730-8000(2008)27%255B21:TROTHV%255D2.0.CO;2

Konn, C., Donval, J. P., Guyader, V., Germain, Y., Alix, A. S., Roussel, E., & Rouxel, O. (2022). Extending the dataset of fluid geochemistry of the Menez Gwen, Lucky Strike, Rainbow, TAG and Snake Pit hydrothermal vent fields: Investigation of temporal stability and organic contribution. Deep-Sea Research Part I: Oceanographic Research Papers, 179(September 2021), 103630. 10.1016/j.dsr.2021.103630

Kopelman, N. M., Mayzel, J., Jakobsson, M., Rosenberg, N. A., & Mayrose, I. (2015). CLUMPAK: A program for identifying clustering modes and packaging population structure inferences across *K*. Molecular Ecology Resources, 15(5), 1179–1191. 10.1111/1755-0998.12387

Krishnan, J., & Rohner, N. (2017). Cavefish and the basis for eye loss. Philosophical Transactions of the Royal Society B: Biological Sciences, 372(1713). 10.1098/rstb.2015.0487

Linse, K., Nye, V., Copley, J. T., & Chen, C. (2019). On the systematics and ecology of two new species of Provanna (Gastropoda: Provannidae) from deep-sea hydrothermal vents in the Caribbean Sea and Southern Ocean. Journal of Molluscan Studies, 85(4), 425–438. 10.1093/mollus/eyz024

Luu, K., Bazin, E., & Blum, M. G. B. (2017). *pcadapt*: An R package to perform genome scans for selection based on principal component analysis. Molecular Ecology Resources, 17(1), 67–77. 10.1111/1755-0998.12592

Matabos, M., Cannat, M., Ballu, V., Barreyre, T., Blandin, J., Castillo, A., Cathalot, C., Chavagnac, V., Chu, N. C., Colaço, A., Crawford, W., Escartin, J., Ferron, B., Fontaine, F., Gautier, L., Godfroy, A., Laes-Huon, A., Lanteri, N., Leau, H.,…Sarradin, P. M. (2025). The EMSO-Azores deep-sea observatory: 15 years of multidisciplinary studies of the lucky strike hydrothermal system, from sub-seafloor to the water column. Journal of Sea Research, 207, 102625. 10.1016/j.seares.2025.102625

Matabos, M., Plouviez, S., Hourdez, S., Desbruyères, D., Legendre, P., Warén, A., Jollivet, D., & Thiébaut, E. (2011). Faunal changes and geographic crypticism indicate the occurrence of a biogeographic transition zone along the southern East Pacific Rise: Biogeographic transition zone along the southern EPR. Journal of Biogeography, 38(3), 575–594. 10.1111/j.1365-2699.2010.02418.x

Menini, E., & Van Dover, C. L. (2019). An atlas of protected hydrothermal vents. Marine Policy, 108, 103654. 10.1016/j.marpol.2019.103654

Methou, P., Guéganton, M., Copley, J. T., Kayama Watanabe, H., Pradillon, F., Cambon-Bonavita, M.-A., & Chen, C. (2024). Distinct development trajectories and symbiosis modes in vent shrimps. Evolution, 78(3), 413–422. 10.1093/evolut/qpad217

Methou, P., Hernández-Ávila, I., Cathalot, C., Cambon-Bonavita, M.-A., & Pradillon, F. (2022). Population structure and environmental niches of Rimicaris shrimps from the Mid-Atlantic Ridge. Marine Ecology Progress Series, 684, 1–20. 10.3354/meps13986

Methou, P., Johnson, S. B., Sherrin, J., Shank, T. M., Chen, C., & Tunnicliffe, V. (2025). A tale of two shrimps – Speciation and demography of two sympatric shrimp species from hydrothermal vents. Molecular Ecology, 34(22). 10.1111/mec.70119

Methou, P., Michel, L. N., Segonzac, M., Cambon-Bonavita, M.-A., & Pradillon, F. (2020). Integrative taxonomy revisits the ontogeny and trophic niches of Rimicaris vent shrimps. Royal Society Open Science, 7(7), 200837. 10.1098/rsos.200837

Methou, P., Ogawa, N. O., Nomaki, H., Ohkouchi, N., Chen, C., & Schnabel, K. (2024). Genetic connectivity and isotopic niches of alvinocaridid shrimps from chemosynthetic habitats in Aotearoa/New Zealand, with a new Alvinocaris species. Marine Ecology Progress Series. 10.3354/meps14611

Mullineaux, L. S., Kim, S. L., Pooley, A., & Lutz, R. A. (1996). Identification of archaeogastropod larvae from a hydrothermal vent community. Marine Biology, 124, 551–560. 10.1007/BF00351036

Mullineaux, L. S., Metaxas, A., Beaulieu, S. E., Bright, M., Gollner, S., Grupe, B. M., Herrera, S., Kellner, J. B., Levin, L. A., Mitarai, S., Neubert, M. G., Thurnherr, A. M., Tunnicliffe, V., Watanabe, H. K., & Won, Y. J. (2018). Exploring the ecology of deep-sea hydrothermal vents in a metacommunity framework. Frontiers in Marine Science, 4(FEB). 10.3389/fmars.2018.00049

Mullineaux, L. S., Mills, S. W., Le Bris, N., Beaulieu, S. E., Sievert, S. M., & Dykman, L. N. (2020). Prolonged recovery time after eruptive disturbance of a deep-sea hydrothermal vent community. Proceedings of the Royal Society B: Biological Sciences, 287(1941), 20202070. 10.1098/rspb.2020.2070

Nye, V., Copley, J., & Plouviez, S. (2012). A new species of *Rimicaris* (Crustacea: Decapoda: Caridea: Alvinocarididae) from hydrothermal vent fields on the Mid-Cayman Spreading Centre, Caribbean. Journal of the Marine Biological Association of the United Kingdom, 92(5), 1057–1072. 10.1017/S0025315411002001

Nye, V., Copley, J. T. P., & Tyler, P. A. (2013). Spatial Variation in the Population Structure and Reproductive Biology of Rimicaris hybisae (Caridea: Alvinocarididae) at Hydrothermal Vents on the Mid-Cayman Spreading Centre. PLoS ONE, 8(3). 10.1371/journal.pone.0060319

O’Mullan, G. D., Maas, P. A. Y., Lutz, R. A., & Vrijenhoek, R. C. (2001). A hybrid zone between hydrothermal vent mussels (Bivalvia: Mytilidae) from the Mid-Atlantic Ridge. Molecular Ecology, 10(12), 2819–2831. 10.1046/j.1365-294X.2001.t01-1-01401.x

Pembleton, L. W., Cogan, N. O. I., & Forster, J. W. (2013). St AMPP: An R package for calculation of genetic differentiation and structure of mixed-ploidy level populations. Molecular Ecology Resources, 13(5), 946–952. 10.1111/1755-0998.12129

Plouviez, S., Jacobson, A., Wu, M., & Van Dover, C. L. (2015). Characterization of vent fauna at the mid-cayman spreading center. Deep-Sea Research Part I: Oceanographic Research Papers, 97, 124–133. 10.1016/j.dsr.2014.11.011

Plouviez, S., Shank, T. M., Faure, B., Daguin-Thiebaut, C., Viard, F., Lallier, F. H., & Jollivet, D. (2009). Comparative phylogeography among hydrothermal vent species along the East Pacific Rise reveals vicariant processes and population expansion in the South. Molecular Ecology, 18(18), 3903–3917. 10.1111/j.1365-294X.2009.04325.x

Poitrimol, C., Thiébaut, E., Daguin-Thiébaut, C., Le Port, A.-S., Ballenghien, M., Tran Lu Y, A., Hourdez, S., & Jollivet, D. (2022). Contrasted phylogeographic patterns of hydrothermal vent gastropods along South West Pacific: Woodlark Basin, a possible contact zone and/or stepping-stone. PLoS ONE, 1–27. 10.1371/journal.pone.0275638

Pond, D. W., Gebruk, A., Southward, E. C., Southward, A. J., Fallick, A. E., Bell, M. V., & Sargent, J. R. (2000). Unusual fatty acid composition of storage lipids in the bresilioid shrimp Rimicaris exoculata couples the photic zone with MAR hydrothermal vent sites. Marine Ecology Progress Series, 198, 171–179. 10.3354/meps198171

Ponsard, J., Cambon-Bonavita, M.-A., Zbinden, M., Lepoint, G., Joassin, A., Corbari, L., Shillito, B., Durand, L., Cueff-Gauchard, V., & Compère, P. (2013). Inorganic carbon fixation by chemosynthetic ectosymbionts and nutritional transfers to the hydrothermal vent host-shrimp Rimicaris exoculata. ISME Journal, 7(1), 96–109. 10.1038/ismej.2012.87

Privé, F., Luu, K., Vilhjálmsson, B. J., & Blum, M. G. B. (2020). Performing Highly Efficient Genome Scans for Local Adaptation with R Package pcadapt Version 4. Molecular Biology and Evolution, 37(7), 2153–2154. 10.1093/molbev/msaa053

Purcell, S., Neale, B., Todd-Brown, K., Thomas, L., Ferreira, M. A. R., Bender, D., Maller, J., Sklar, P., De Bakker, P. I. W., Daly, M. J., & Sham, P. C. (2007). PLINK: A Tool Set for Whole-Genome Association and Population-Based Linkage Analyses. The American Journal of Human Genetics, 81(3), 559–575. 10.1086/519795

Rochette, N. C., & Catchen, J. M. (2017). Deriving genotypes from RAD-seq short-read data using Stacks. Nature Protocols, 12(12), 2640–2659. 10.1038/nprot.2017.123

Rochette, N. C., Rivera-Colón, A. G., & Catchen, J. M. (2019). Stacks 2: Analytical methods for paired-end sequencing improve RADseq-based population genomics. Molecular Ecology, 28(21), 4737–4754. 10.1111/mec.15253

Roux, C., Fraïsse, C., Romiguier, J., Anciaux, Y., Galtier, N., & Bierne, N. (2016). Shedding Light on the Grey Zone of Speciation along a Continuum of Genomic Divergence. PLOS Biology, 14(12), e2000234. 10.1371/journal.pbio.2000234

Sarrazin, J., Cathalot, C., Laes, A., Marticorena, J., Michel, L. N., & Matabos, M. (2022). Integrated Study of New Faunal Assemblages Dominated by Gastropods at Three Vent Fields Along the Mid-Atlantic Ridge: Diversity, Structure, Composition and Trophic Interactions. Frontiers in Marine Science, 9, 925419. 10.3389/fmars.2022.925419

Sarrazin, J., Portail, M., Legrand, E., Cathalot, C., Laes, A., Lahaye, N., Sarradin, P. M., & Husson, B. (2020). Endogenous versus exogenous factors: What matters for vent mussel communities? Deep Sea Research Part I: Oceanographic Research Papers, 160, 103260. 10.1016/j.dsr.2020.103260

Schwander, T., Lo, N., Beekman, M., Oldroyd, B. P., & Keller, L. (2010). Nature versus nurture in social insect caste differentiation. Trends in Ecology and Evolution, 25(5), 275–282. 10.1016/j.tree.2009.12.001

Segonzac, M., de Saint Laurent, M., & Casanova, B. (1993). L’enigme du comportement trophique des crevettes Alvinocarididae des sites hydrothermaux de la dorsale medio-atlantique. Cahiers de Biologie Marine, 34(4), 535–571. 10.21411/%2520CBM.A.B3683E29

Streit, K., Bennett, S. A., Van Dover, C. L., & Coleman, M. (2015). Sources of organic carbon for *Rimicaris hybisae*: Tracing individual fatty acids at two hydrothermal vent fields in the Mid-Cayman rise. Deep Sea Research Part I: Oceanographic Research Papers, 100, 13–20. 10.1016/j.dsr.2015.02.003

Sundqvist, L., Keenan, K., Zackrisson, M., Prodöhl, P., & Kleinhans, D. (2016). Directional genetic differentiation and relative migration. Ecology and Evolution, 6(11), 3461–3475. 10.1002/ece3.2096

Teixeira, S., Cambon-Bonavita, M. A., Serrão, E. A., Desbruyéres, D., & Arnaud-Haond, S. (2011). Recent population expansion and connectivity in the hydrothermal shrimp Rimicaris exoculata along the Mid-Atlantic Ridge. Journal of Biogeography, 38(3), 564–574. 10.1111/j.1365-2699.2010.02408.x

Teixeira, S., Olu, K., Decker, C., Cunha, R. L., Fuchs, S., Hourdez, S., Serrão, E. A., & Arnaud-Haond, S. (2013). High connectivity across the fragmented chemosynthetic ecosystems of the deep Atlantic Equatorial Belt: Efficient dispersal mechanisms or questionable endemism? Molecular Ecology, 22(18), 4663–4680. 10.1111/mec.12419

Teixeira, S., Serrão, E. A., & Arnaud-Haond, S. (2012). Panmixia in a fragmented and unstable environment: The hydrothermal shrimp Rimicaris exoculata disperses extensively along the Mid-Atlantic ridge. PLoS ONE, 7(6), 1–10. 10.1371/journal.pone.0038521

Thaler, A. D., Plouviez, S., Saleu, W., Alei, F., Jacobson, A., Boyle, E. A., Schultz, T. F., Carlsson, J., & Van Dover, C. L. (2014). Comparative population structure of two deep-sea hydrothermal-vent-associated decapods (Chorocaris sp. 2 and Munidopsis lauensis) from southwestern Pacific back-arc basins. PLoS ONE, 9(7), 1–13. 10.1371/journal.pone.0101345

Tran Lu Y, A., Ruault, S., Daguin-Thiébaut, C., Castel, J., Bierne, N., Broquet, T., Wincker, P., Perdereau, A., Arnaud-Haond, S., Gagnaire, P.-A., Jollivet, D., Hourdez, S., & Bonhomme, F. (2022). Subtle limits to connectivity revealed by outlier loci within two divergent metapopulations of the deep-sea hydrothermal gastropod Ifremeria nautilei. Molecular Ecology, 1–46. 10.1111/mec.16430

Tunnicliffe, V., & Fowler, C. M. R. (1996). Influence of sea-floor spreading on the global hydrothermal vent fauna. Letters to Nature, 379. 10.1038/379531a0

Tunnicliffe, V., St. Germain, C., & Hilário, A. (2014). Phenotypic Variation and Fitness in a Metapopulation of Tubeworms (Ridgeia piscesae Jones) at Hydrothermal Vents. PLoS ONE, 9(10), e110578. 10.1371/journal.pone.0110578

Tyler, P. A., & Dixon, D. R. (2000). Temperature/pressure tolerance of the first larval stage of Mirocaris fortunata from Lucky Strike hydrothermal vent field. Journal of the Marine Biological Association of the United Kingdom, 80(4), 739–740. 10.1017/S0025315400002605

Van Audenhaege, L., Sarrazin, J., Legendre, P., Perrois, G., Cannat, M., Arnaubec, A., & Matabos, M. (2024). Monitoring ecological dynamics on complex hydrothermal structures: A novel photogrammetry approach reveals fine-scale variability of vent assemblages. Limnology and Oceanography, 69(2), 325–338. 10.1002/lno.12486

van der Heijden, K., Petersen, J. M., Dubilier, N., & Borowski, C. (2012). Genetic connectivity between north and south Mid-Atlantic Ridge chemosynthetic bivalves and their symbionts. PLoS ONE, 7(7). 10.1371/journal.pone.0039994

Van Der Heijden, K., Petersen, J. M., Dubilier, N., & Borowski, C. (2012). Genetic Connectivity between North and South Mid-Atlantic Ridge Chemosynthetic Bivalves and Their Symbionts. PLoS ONE, 7(7), e39994. 10.1371/journal.pone.0039994

Vereshchaka, A. L., Kulagin, D. N., & Lunina, A. A. (2015). Phylogeny and new classification of hydrothermal vent and seep shrimps of the family alvinocarididae (Decapoda). PLoS ONE, 10(7), 1–29. 10.1371/journal.pone.0129975

Versteegh, E. A. A., Van Dover, C. L., Van Audenhaege, L., & Coleman, M. (2023). Multiple nutritional strategies of hydrothermal vent shrimp (*Rimicaris hybisae*) assemblages at the Mid-Cayman Rise. Deep Sea Research Part I: Oceanographic Research Papers, 192, 103915. 10.1016/j.dsr.2022.103915

Weir, B. S., & Cockerham, C. C. (1984). Estimating F-Statistics for the Analysis of Population Structure. Evolution, 38(6), 1358–1370. 10.2307/2408641

Yahagi, T., Fukumori, H., Warén, A., & Kano, Y. (2019). Population connectivity of hydrothermal-vent limpets along the northern Mid-Atlantic Ridge (Gastropoda: Neritimorpha: Phenacolepadidae). Journal of the Marine Biological Association of the United Kingdom, 99(1), 179–185. 10.1017/S0025315417001898

Yahagi, T., Kayama Watanabe, H., Kojima, S., Kano, Y., Watanabe, H. K., Kojima, S., Kano, Y., Kayama Watanabe, H., Kojima, S., & Kano, Y. (2017). Do larvae from deep-sea hydrothermal vents disperse in surface waters? Ecology, 98(6), 1524–1534. 10.1002/ecy.1800

Yahagi, T., Watanabe, H., Ishibashi, J. I., & Kojima, S. (2015). Genetic population structure of four hydrothermal vent shrimp species (Alvinocarididae) in the Okinawa Trough, Northwest Pacific. Marine Ecology Progress Series, 529, 159–169. 10.3354/meps11267

Zbinden, M., & Cambon-Bonavita, M. (2020). Biology and ecology of Rimicaris exoculata, a symbiotic shrimp from deep-sea hydrothermal vents. Marine Ecology Progress Series, 652, 187–222. 10.3354/meps13467

Zhou, H., Alexander, D., & Lange, K. (2011). A quasi-Newton acceleration for high-dimensional optimization algorithms. Statistics and Computing, 21(2), 261–273. 10.1007/s11222-009-9166-3

Zhou, Y., Chen, C., Zhang, D., Wang, Y., Kayama, H., Jin, W., Dass, S., Ruiyan, B., Han, Y., Sun, D., Xu, P., Lu, B., Zhai, H., Han, X., Tao, C., Qiu, Z., Sun, Y., Liu, Z., Qiu, W., & Wang, C. (2022). Delineating biogeographic regions in Indian Ocean deep-sea vents and implications for conservation. Diversity and Distributions, April, 1–13. 10.1111/ddi.13535

