## Supplementary Methods & Figures S1 to S8 for "High gene flow in Atlantic vent shrimp species with multiple ridge colonizations by ecotypic lineages with contrasting symbioses"

**Deep-sea Atlantic vent shrimp species display widespread gene flow and multiple ridge colonization despite ecotypic lineages with contrasting symbioses.**

Portanier Elodie^1,2§^, Methou Pierre^1§*^, Daguin-Thiébaut Claire^2^, Ruault Stéphanie^2^, Le Port Anne-Sophie^2^, Chen Chong^4^, Matabos Marjolaine^1^, Jollivet Didier ^2^, Pradillon Florence^1*^,

^1^Univ Brest, Ifremer, BEEP, F-29280 Plouzané, France.

^2^Sorbonne Université, CNRS, UMR 7144 ‘AD2M’, DISEEM lab, Station Biologique de Roscoff, Place Georges Teissier, Roscoff, France.

^3^Institut de Systématique, Evolution, Biodiversité (ISYEB), Muséum national d'Histoire naturelle, CNRS, SU, EPHE-PSL, UA, 57 rue Cuvier, CP51, 75005 Paris, France

^4^X-STAR, Japan Agency for Marine-Earth Science and Technology (JAMSTEC), 2-15 Natsushima-cho, Yokosuka, Kanagawa 237-0061, Japan

**Details of Stacks parameters optimisation and SNP filtering**

We followed the r0.80 procedure detailed in Paris et al. (2017) and Rochette and Catchen (2017) to optimize Stacks parameters. The *denovo_map.pl* function was thus ran on a subset of 24 (*R. exoculata*) or 40 (*R. chacei*, *R. hybisae*) samples representing all vent localities and selected to have the maximal number of reads in each locality, to generate a locus catalogue. Values from 1 to 9 were tested for the number of mismatches allowed between reads to create a stack within individuals (*-M*) and the number of mismatches allowed between individuals to group stacks (*-n*) with *M* = *n*. The minimum number of reads required to generate an initial stack (*-m*) was fixed to 3. The minimum percentage of individuals that share a RAD locus for a given population in the *populations* module was set to *-r* = 0.80. The optimal values for *-M* = *-n* were identified as the ones from which the number of polymorphic loci, the number of SNPs and the percentage of loci showing specific number of SNPs started to plateau. The selected values were -M and -n = 4 and 6 for *R. exoculata* and for *R. chacei / R. hybisae*, respectively (see Supplementary Figures S1 and S2).

Data were subsequently filtered using VCFtools (v.0.1.16) (Danecek et al. 2011) and R v.4.1.0 (R Core Team 2021). Only biallelic SNPs with a minimum coverage of 10 reads and a maximum coverage not greater than third the average read coverage across nucleotide sites (i.e. SNPs with coverage > 165 and 210 for *R. exoculata* and *R. chacei / R. hybisae*, respectively) were kept. Maximum coverage filter intends to eliminate potential unfiltered paralogs.

Triplicated individuals were used to identify and exclude SNPs that were erroneously genotyped along the genome as follows: if the same SNP was genotyped differently between replicated individuals, it was excluded from the dataset for all individuals. Among the three replicates for each individual, the one showing the lowest percentage of missing data was then kept in the final dataset. Finally, SNPs and individuals with more than 10% of missing data were excluded.

To avoid the presence of SNPs in linkage disequilibrium, a single SNP per locus was kept and outlier SNPs were excluded using the R package pcadapt v. 4.3.3 (Luu et al. 2017, Privé et al. 2020). For each species, two principal components (PCs) were used to eliminate outliers (see Supplementary Figures S3 and S4) following the Cattell’s rule (i.e. the optimal number of PCs is given from the break line in the scree-plot, with the following PCs accounting only for random variation). All SNPs showing p-values < 0.05, after adjustment using q-values, a false discovery rate (Benjamini and Hochberg 1995) or Bonferroni correction, were excluded from further analyses.

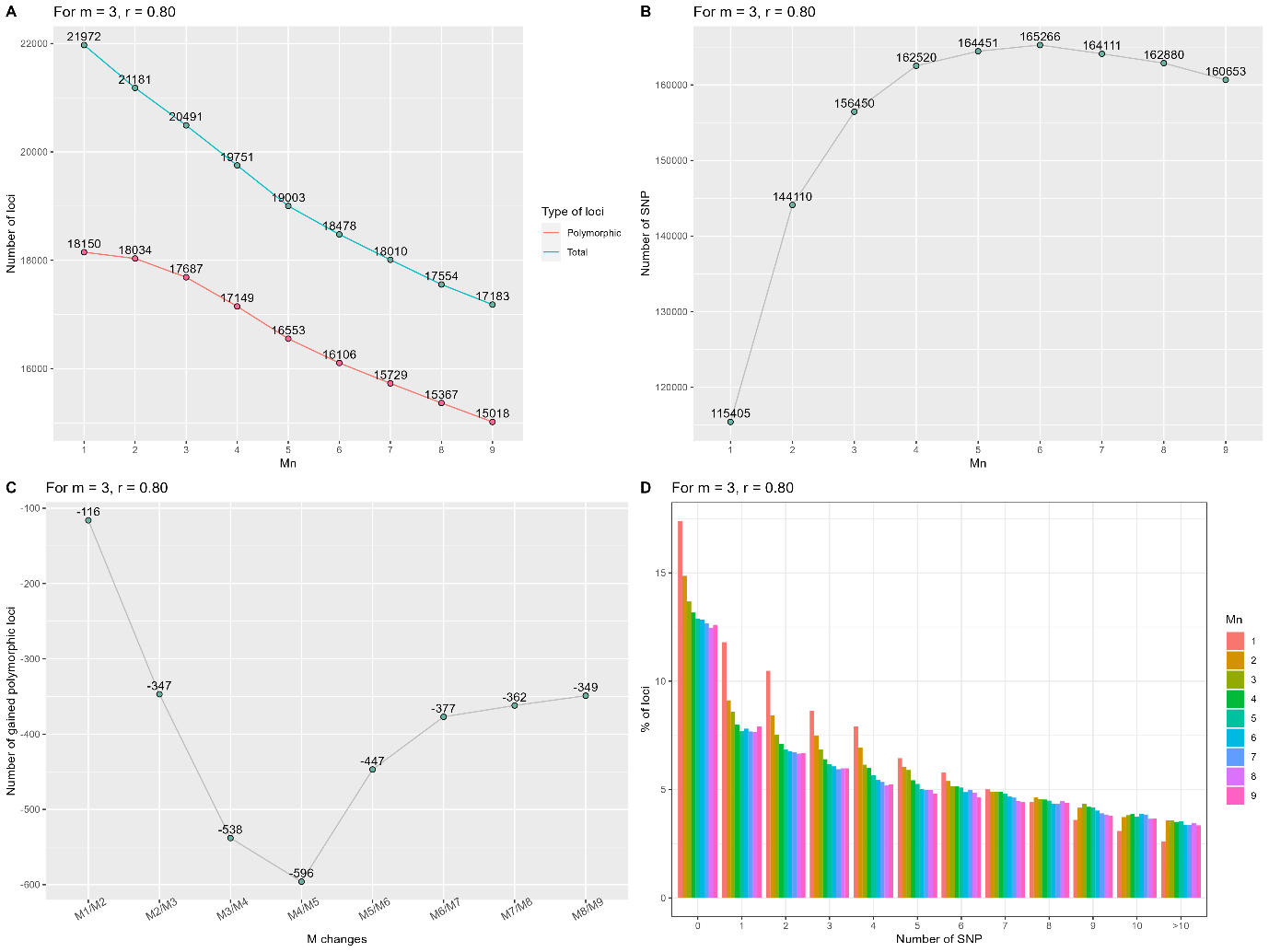

Figure S1: Stacks parameters optimization for *Rimicaris exoculata*. A) Evolution of the total number of assembled de novo loci and of the number of polymorphic loci among those according to *M* and *n* parameters values. B) Evolution of the total number of SNPs (several per loci) according to *M* and *n* parameters values. C) Number of gained (or lost if negative) polymorphic loci between two *M* and *n* parameters values. D) Percentage of loci showing specific number of SNPs according to *M* and *n* parameters values.

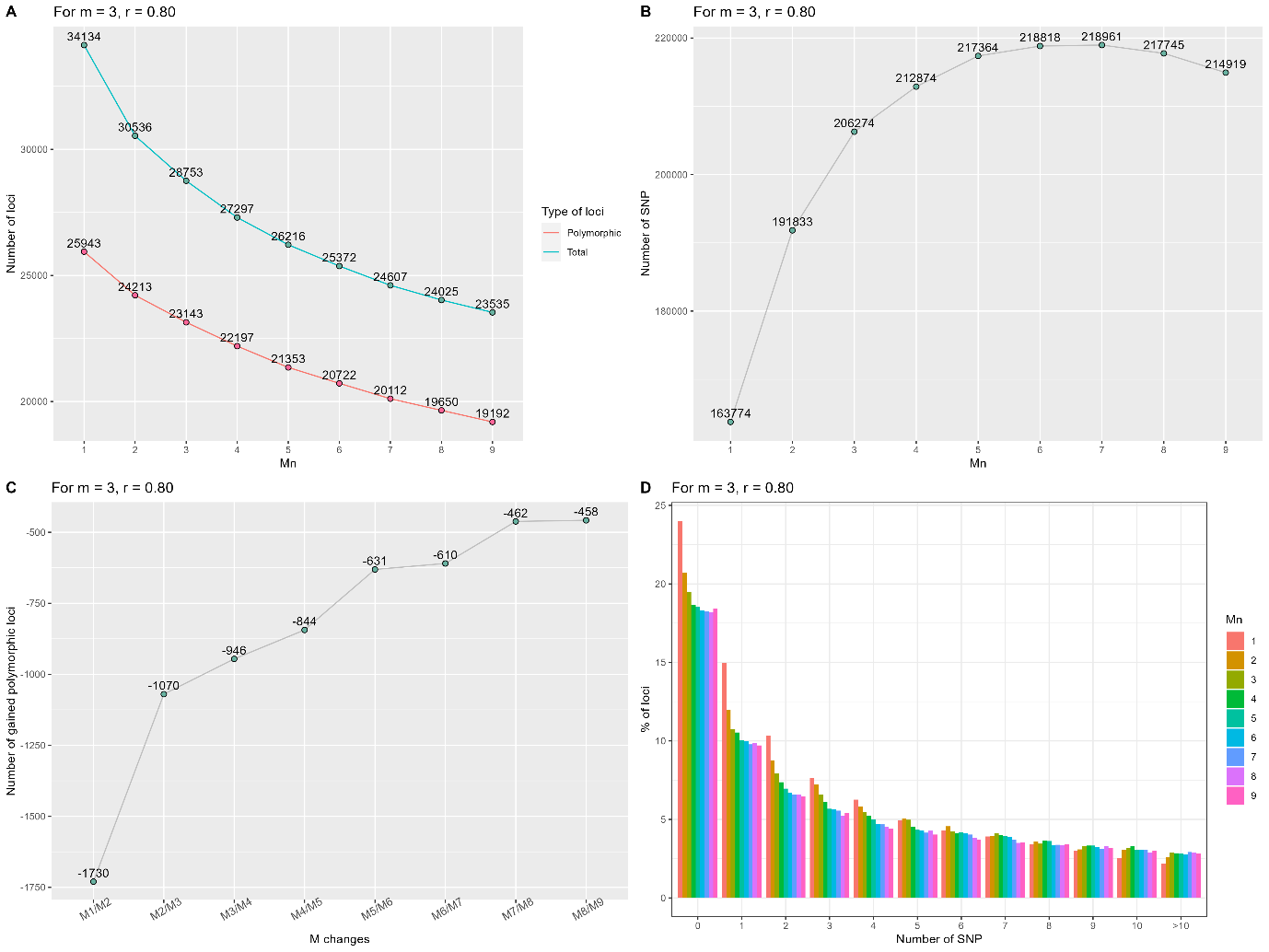

Figure S2: Stacks parameters optimization for *Rimicaris chacei / R. hybisae* assembly. A) Evolution of the total number of assembled de novo loci and of the number of polymorphic loci among those according to *M* and *n* parameters values. B) Evolution of the total number of SNPs (several per loci) according to *M* and *n* parameters values. C) Number of gained (or lost if negative) polymorphic loci between two *M* and *n* parameters values. D) Percentage of loci showing specific number of SNPs according to *M* and *n* parameters values.

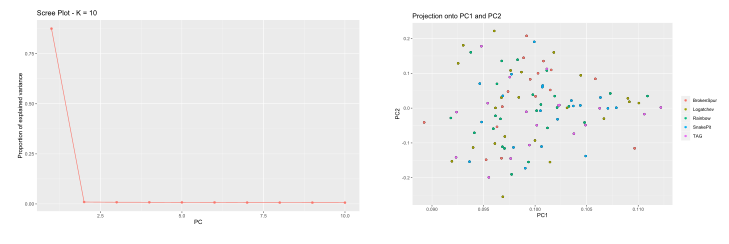

Figure S3: Pcadapt scree-plot and projections used to choose the number of PCs to keep in outlier detection analysis performed for *R. exoculata*.

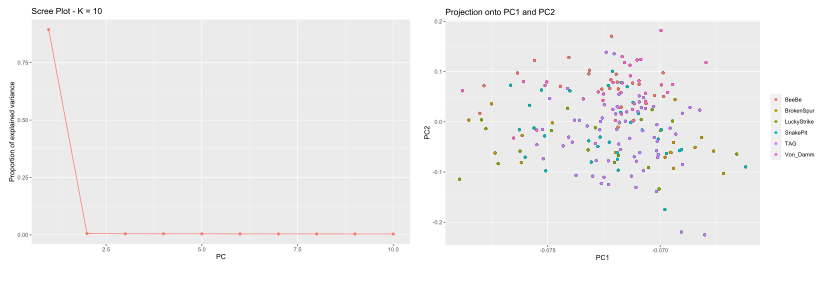

Figure S4: Pcadapt scree-plot and projections used to choose the number of PCs to keep in outlier detection analysis performed for *R. chacei / R. hybisae*.

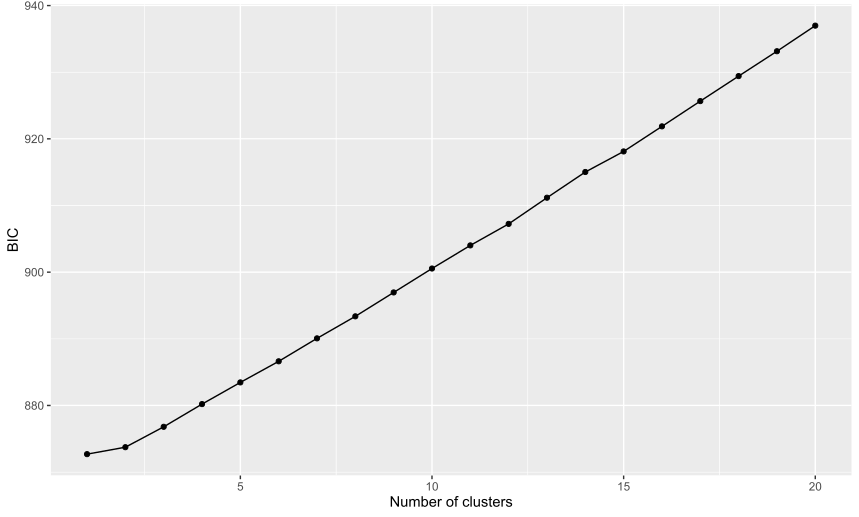

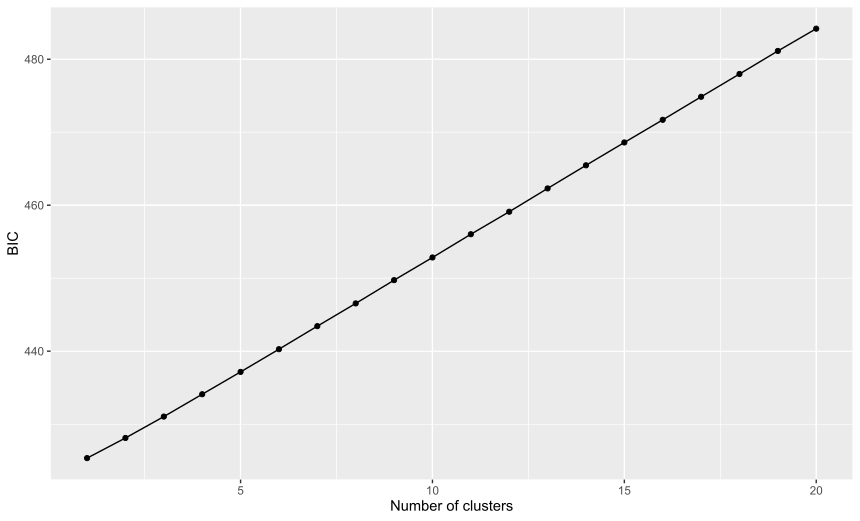

Figure S5: BIC values from the DAPC procedure for *R. chacei / R. hybisae* (top) and *R. exoculata* (bottom), sampled along the Mid-Atlantic Ridge and in the Mid-Cayman spreading center.

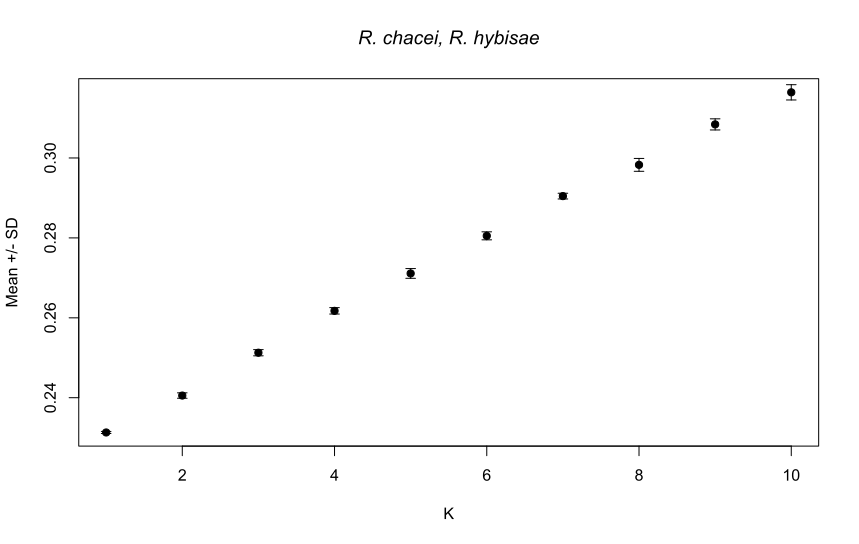

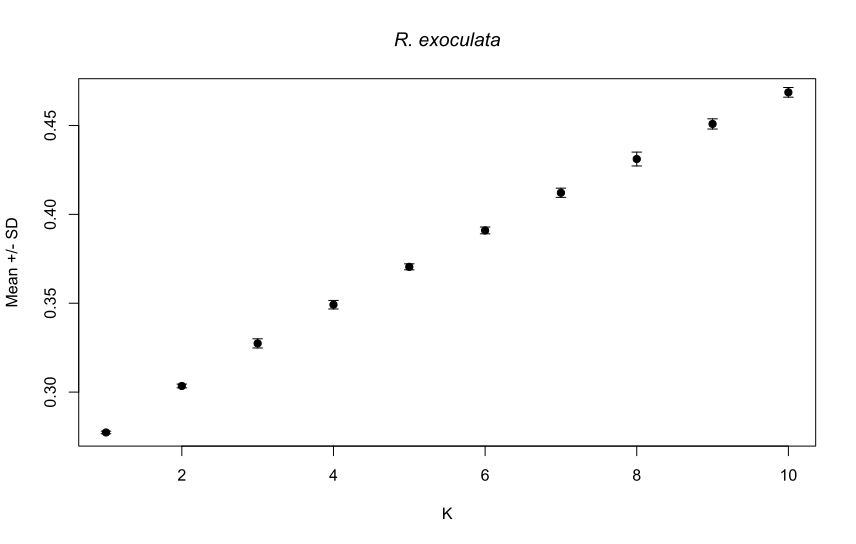

Figure S6: Average CV-errors values across the 10 runs of ADMIXTURE for *R. chacei/R. hybisae* and *R. exoculata* assemblies.

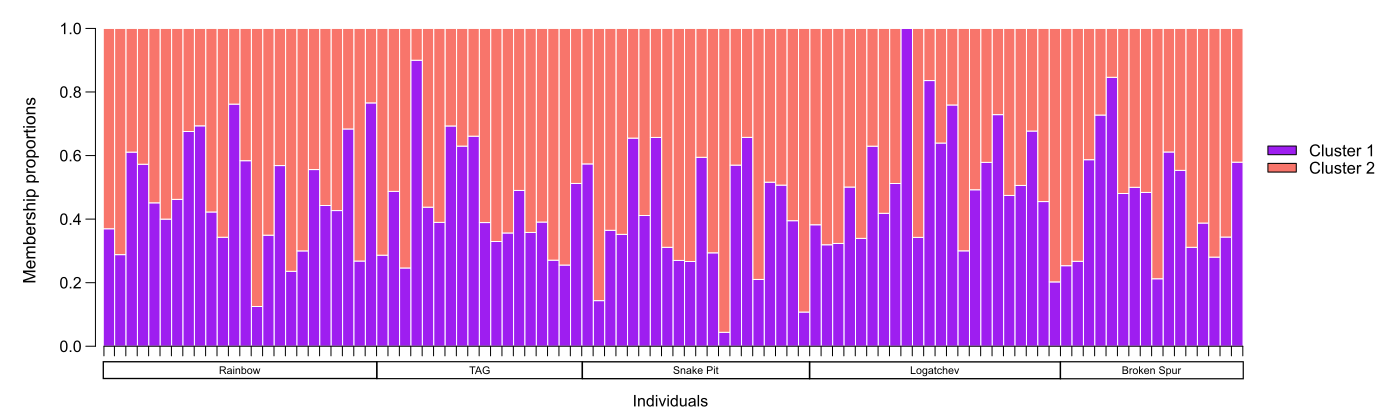

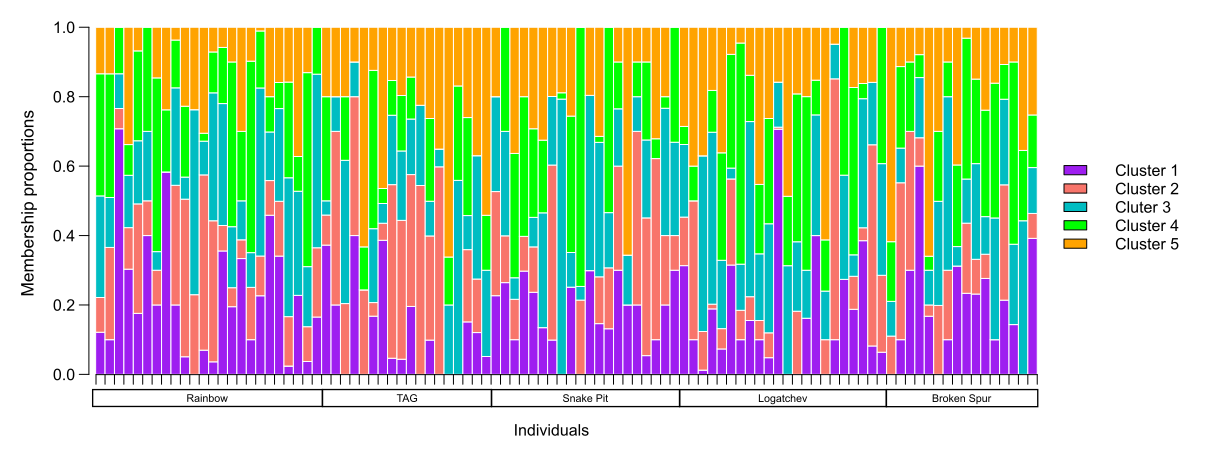

Figure S7: ADMIXTURE membership proportions for *R. exoculata* from the Mid-Atlantic Ridge when considering K = 2 or K = 5 genetic clusters.

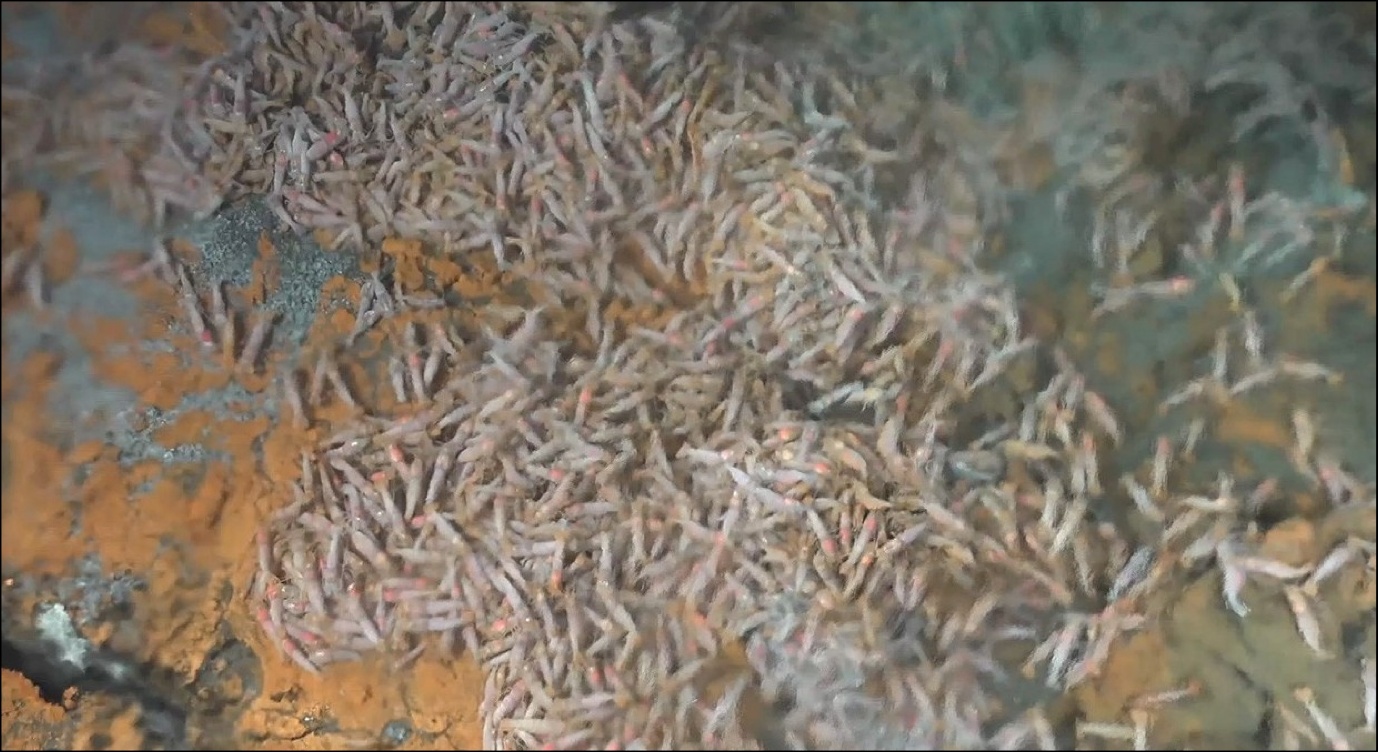

Figure S8: A rare swarm of *R. chacei* adults near shimmering vent fluids recently discovered at the TAG vent field in November 2023 (Bicose 3 expedition).

**References**

Benjamini, Y. & Hochberg, Y. (1995) Controlling the False Discovery Rate: a practical and powerful approach to multiple testing. Journal of the Royal Statistical Society: Series B (Methodological), 57: 289-300.

Danecek P., Auton A., Abecasis G., Albers, C.A., Banks E., DePristo M.A., Handsaker R., Lunter, G., Marth, G., Sherry, S.T., McVean G., Durbin, R., 1000 Genomes Project Analysis Group. (2011) The Variant Call Format and VCFtools. Bioinformatics, 27, 2156–2158.

Luu, K., Bazin, E., & Blum, M.G.B. (2017). pcadapt: An R package to perform genome scans for selection based on principal component analysis. Molecular Ecology Resources, 17, 67–77.

Paris, J. R., Stevens, J. R., & Catchen, J. M. (2017). Lost in parameter space: A road map for stacks. Methods in Ecology and Evolution, 8, 1360‑1 7.

Privé, F., Luu, K., Vilhjálmsson, B. J., & Blum, M. G.B. (2020) Performing highly efficient genome scans for local adaptation with R package pcadapt version 4. Molecular Biology and Evolution, 37, 2153–2154.

Rochette, N.C., & Catchen, J.M. (2017) Deriving genotypes from RAD-seq short-read data using Stacks. Nature Protocols, 12, 2640–2659.
